# Serine/arginine-rich splicing factor 7 plays oncogenic roles through specific regulation of m^6^A RNA modification

**DOI:** 10.1101/2021.10.11.463901

**Authors:** Yixian Cun, Sanqi An, Haiqing Zheng, Jing Lan, Wenfang Chen, Wanjun Luo, Chengguo Yao, Xincheng Li, Xiang Huang, Xiang Sun, Zehong Wu, Yameng Hu, Ziwen Li, Shuxia Zhang, Geyan Wu, Meisongzhu Yang, Miaoling Tang, Ruyuan Yu, Xinyi Liao, Guicheng Gao, Wei Zhao, Jinkai Wang, Jun Li

## Abstract

Serine/Arginine-Rich Splicing Factor 7 (SRSF7), which is previously recognized as a splicing factor, has been revealed to play oncogenic roles in multiple cancers. However, the mechanisms underlying its oncogenic roles have not been well addressed. Here, based on N6-methyladenosine (m^6^A) co-methylation network analysis across diverse cell lines, we found SRSF7 positively correlated with glioblastoma cell-specific m^6^A methylation. We then proved SRSF7 is a novel m^6^A regulator that specifically facilitates the m^6^A methylation near its binding sites on the mRNAs involved in cell proliferation and migration through recruiting methyltransferase complex. Moreover, SRSF7 promotes the proliferation and migration of glioblastoma cells largely dependent on the m^6^A methyltransferase. The two single-nucleotide m^6^A sites on *PBK* are regulated by SRSF7 and partially mediate the effects of SRSF7 on glioblastoma cells through recognition by IGF2BP2. Together, our discovery revealed a novel role of SRSF7 in regulating m^6^A and timely confirmed the existence and functional importance of RNA binding protein (RBP) mediated specific regulation of m^6^A.

## Introduction

Serine/arginine-rich splicing factor 7 (SRSF7, also known as 9G8) belongs to the serine/arginine (SR) protein family, which contains 7 canonical members (SRSF1-7) [1]. It is previously known as a splicing factor to regulate alternative splicing as well as a regulator of alternative polyadenylation [2–5]. SRSF7 is also an adaptor of NXF1, which exports mature RNAs out of nucleus, and plays important roles in coupling RNA alternative splicing and polyadenylation to mRNA export [5]. It was reported that hyperphosphorylated SRSF7 binds to pre-mRNA for splicing and it becomes hypophosphorylated during splicing, the later form of SRSF7 can bind the NXF1 for the subsequent export of the spliced RNAs [3].

The oncogenic roles of SRSF7 have been widely reported. It was discovered as a critical gene required for cell growth or viability in multiple cancer cell lines based on a genome-wide CRISPR-Cas9 screening [6]. Aberrantly elevated expression of SRSF7 had been observed in lung cancer, colon cancer and gastric cancer [7–9]. It was also reported to be highly expressed in glioblastoma (GBM, grade IV glioma) and associated with poor patient outcome [10]. However, although SRSF7 has been reported to regulate splicing, APA, and mRNA export, the mechanisms underlying its oncogenic roles have not been well addressed.

N6-methyladenosine (m^6^A) is a reversible RNA modification prevalent in eukaryotic messenger RNAs (mRNAs) and long non-coding RNA [11–13]. It plays critical roles in various biological process, including stem cell differentiation, immune system, learning and memory, cancer development [14–18]. The m^6^A modification is marked by the m^6^A methyltransferase (also known as “writers”) complex, which consists of METTL3, METTL14, WTAP, VIRMA, ZC3H13, RBM15/15B and HAKAI (also known as CBLL1) [19, 20]. m^6^A can also be removed by demethylases (also known as “erasers”) including FTO and ALKBH5 [21, 22]. The m^6^A-modified RNAs are recognized by a series of readers such as YTH-domain containing proteins (YTHDF1-3 and YTHDC1-2), in which YTHDF2 facilitates the degradation of methylated RNAs and is important for cell fate transitions [23–26]. IGF2BP1-3 are a different type of readers that can stabilize the methylated RNAs and play oncogenic roles in multiple types of cancers [27]. In addition, m^6^A can also down-regulate gene expression through degrading chromosome-associated regulatory RNAs (carRNAs) [28] and up-regulate gene expression by demethylating H3K9me2 histone modification [29].

Unlike global regulation of m^6^A by the methyltransferase complex, selective modification of m^6^A on specific targets can shape the cell-specific methylome and mediate specific functions in diverse biological systems. There are different mechanisms that confer the specificities of m^6^A. Although the components of methyltransferase complex VIRMA and ZC3H13 mainly affect the m^6^A at stop codon and 3’ UTR, their substantial effects on m^6^A suggest fundamental but limited specificities for m^6^A installation, consistent with that they do not have RNA binding domain and ZC3H13 works to take the methyltransferase into nucleus [30, 31]. Since m^6^A occurs co-transcriptionally, m^6^A could be specifically regulated co-transcriptionally through H3K36me3 and transcription factors. Depletion of H3K36me3 also resulted in global reduction of m^6^A, especially the m^6^A at 3’UTRs and protein-coding regions, suggesting a fundamental but relatively low specificity in regulation of m^6^A [32]. On the other hand, transcription factors CEBPZ and SMAD2/3 can recruit the methyltransferase to methylate the nascent RNAs being transcribed by them and play important roles in acute myeloid leukemia oncogenesis and stem cells differentiation respectively [33]. The specificities of transcription factors are conferred by their binding specificities on the promoters. Therefore, they can mediate highly specific methylation other than global regulation of m^6^A. However, transcription factors usually bind at the 5’ end thus cannot precisely direct the m^6^A modification at specific loci of the RNAs. In contrast to transcription factors, which select RNAs other than sites, RBPs have the potential to precisely guide the methylation at specific sites of RNAs in the similar manner as they regulate alternative splicing [34]. Recently, we developed a co-methylation network based computational framework and revealed a large number of RBPs act as m^6^A trans-regulators to specifically regulate m^6^A to form cell-specific m^6^A methylomes [35]. However, firm experimental validations and profound characterizations are still lacking, and whether these RBPs play important functional roles through regulating the m^6^A of specific sites are not clear either.

In this study, we found SRSF7 specifically regulates the m^6^A on genes involved in cell proliferation and migration and plays oncogenic roles through recruiting the m^6^A methyltransferase near its binding sites in GBM cells. Our discovery revealed a novel role of SRSF7 in regulating m^6^A and timely confirmed the existence and importance of RBP-mediated specific regulation of m^6^A.

## Results

### SRSF7 is a potential m^6^A regulator that interacts with m^6^A methyltransferase complex

To elucidate how cells establish cell-specific m^6^A methylomes, we previously developed a co-methylation network based computational framework to systematically identify the cell-specific *trans*-regulators of m^6^A [35]. We first identified the RBPs with gene expression correlated with the m^6^A ratio (level) of specific co-methylation module (a subset of co-methylated m^6^A peaks) across 25 different cell lines (the detailed information of cell lines can be found in the supplementary table of [35]). By further investigating the enrichment of binding targets of the RBPs within their correlated modules based on CLIP-seq data of 157 RBPs and motifs of 89 RBPs, we revealed widespread cell-specific trans-regulation of m^6^A and predicted 32 high-confidence m^6^A regulators [35]. It is of great importance to understand whether these RBP-mediated specific regulation of m^6^A plays critical functional roles. This co-methylation network provides the information about cell specificities of different modules, which give valuable clues for us to speculate the functions of these modules. We realized that one of the modules (M5) were highly methylated in two glioblastoma (GBM) cell lines (PBT003 and GSC) (Figure 1A). Coincidently, although not significant enough to bear multiple testing correction, the mostly enriched Gene Ontology (GO) terms for the corresponding genes of this module are glioma and cancer related pathways, suggesting that the specific methylation of this module may play a role in the development of glioma (Figure 1B). We then tried to dissect the RBPs that direct the specific m^6^A methylation of this glioma related module. As we have previously determined [35] and shown at the bottom of Figure 1A, there were 6 RBPs with gene expression significantly correlated with the m^6^A index (the first component of PCA) of module M5, including 2 positive and 4 negative correlations. We further analyzed the prognostic relevance of these 6 RBPs in GBM patients from Chinese Glioma Genome Atlas (CGGA) dataset [36]. We found that the expression of *SRSF7* was most significantly correlated with the survival time of GBM patients (Figure 1C). Highly expression of *SRSF7* was associated with highly m^6^A methylation of the m^6^A sites in this module and poor prognosis of the GBM patients (Figure 1D and E). Although the other 5 RBPs may also regulate m^6^A of this module in GBM cells, they cannot really affect the prognosis of GBM patients, we therefore focused on SRSF7 to investigate whether and how it plays important role in GBM through specific regulation of m^6^A.

**Figure 1.**
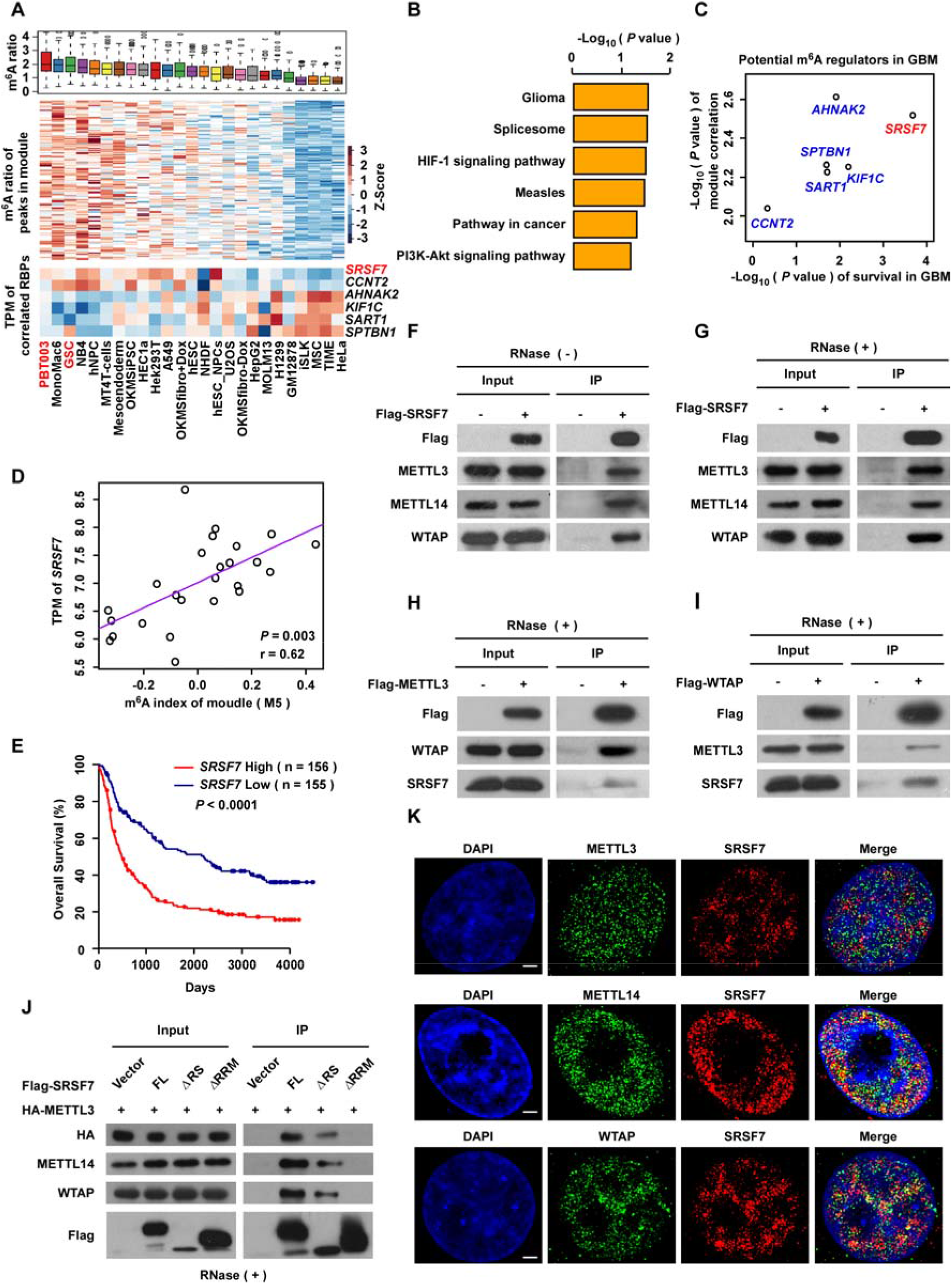
SRSF7 is a potential m^6^A regulator that interacts with m^6^A methyltransferase complex. **A.** The boxplot (upper panel) and heatmap representing the m^6^A ratios of the m^6^A peaks within the co-methylation module M5 as well as the heatmap representing the gene expressions of the RBPs that significantly correlated with the m^6^A indexes of M5 (lower panel). The cell lines are sorted according to the m^6^A indexes of M5, glioblastoma cell lines were colored red. **B.** GO enrichment analysis of corresponding genes in module M5. **C.** The x-axis represents the logarithm transformed *P* values of the correlations between the expression of RBPs and the m^6^A indexes of co-methylation module M5; the y-axis represents the logarithm transformed *P* values of the overall survival (OS) of these 6 RBPs in GBM patients. **D.** Scatter plots representing the correlation between the expression of *SRSF7* and m^6^A indexes of module M5 across 25 cell lines. The *P* value and correlation coefficient are indicated at the bottom right corner. **E.** Kaplan-Meier analysis of overall survival (OS) based on *SRSF7* expression of GBM patients from CGGA dataset. **F, G.** Western blots showing Flag-tagged SRSF7 interacts with endogenous METTL3, METTL14 and WTAP without (F) and with (G) RNase treatment respectively in U87MG cells. **H, I.** Western blots showing Flag-tagged METTL3 (H) and WTAP (I) interact with endogenous SRSF7 with RNase treatment in U87MG cells. **J.** Western blots showing Flag-tagged full-length and truncated SRSF7 interact with HA-tagged METTL3 and endogenous METTL14 and WTAP with RNase treatment in U87MG cells. **K.** 3D-SIM imaging indicating SRSF7 is co-localized with METTL3, METTL14, and WTAP in the nucleus, Scale Bar: 2 μm.

To test whether SRSF7 is a genuine m^6^A regulator that facilitates the installation of m^6^A at specific m^6^A sties, we first examined whether SRSF7 can interact with the core m^6^A methyltransferase complex composed of METTL3, METTL14, and WTAP in a GBM cell line U87MG. Co-immunoprecipitation (Co-IP) assays revealed that Flag-tagged SRSF7 could pull down the endogenous METTL3, METTL14, and WTAP independent of RNA (Figure 1F and G). Reciprocally, both Flag-tagged METTL3 and WTAP could also pull down endogenous SRSF7 in an RNA independent manner respectively in U87MG cells (Figure 1H and I).

Similar results were observed in 293T cells, suggesting the interaction between SRSF7 and methyltransferase complex is a universal mechanism (Figure S1A). In addition, we performed co-IP using truncated SRSF7 with RRM (RNA recognition motif) domain and RS (arginine/serine) domain deleted respectively in U87MG cells and found deletion of RRM domain other than RS domain could disrupt the interaction with METTL3, METTL14, and WTAP, indicating that SRSF7 interacts with the methyltransferase complex through its RPM domain (Figure 1J and Figure S1B).

We then used 3D-SIM super-resolution microscopy to test the protein colocalization between SRSF7 and m^6^A methyltransferase complex in U87MG cells. We found a portion of SRSF7 proteins were colocalized with portions of METTL3, METTL14, and WTAP in the nuclear respectively, implying that at least a part of SRSF7 proteins can specifically regulate m^6^A (Figure 1K). The above results suggest that SRSF7 may be able to regulate m^6^A through recruiting the m^6^A methyltransferase complex.

### SRSF7 specifically facilitates the m^6^A modification near its binding sites

To further investigate whether SRSF7 regulates m^6^A modification, we knocked down SRSF7 and performed m^6^A-seq to examine the m^6^A alteration due to *SRSF7* depletion in U87MG cells. The typical m^6^A motif was enriched in the m^6^A peaks of both knockdown and control cells (Figure S2A). As shown in Figure S2B, the m^6^A peaks were enriched near the stop codons in both knockdown and control cells, which is consistent with previous studies [11, 12]. In contrast to the RBPs in the m^6^A methyltransferase complex, which usually cause massive loss of m^6^A upon depletion [20], depletion of *SRSF7* did not alter the distribution (Figure S2B) and overall peak intensities of the m^6^A peaks (Figure S2C), suggesting that SRSF7 may be a different type of m^6^A regulator that regulates a small number of highly specific m^6^A sites in U87MG cells.

We then determined the differentially methylated m^6^A sites between *SRSF7* knockdown and control to understand the specific sites regulated by SRSF7. After *SRSF7* knockdown, 3334 m^6^A peaks in 2440 genes were down-regulated; in contrast, only 2447 peaks in 1850 genes were up-regulated (Figure 2A and Figure S2D). Gene ontology (GO) analysis and KEGG analysis showed that these differentially methylated genes were enriched in terms including cell division, cell migration, cell proliferation, and pathway in cancer (Figure S2E and F).

**Figure 2.**
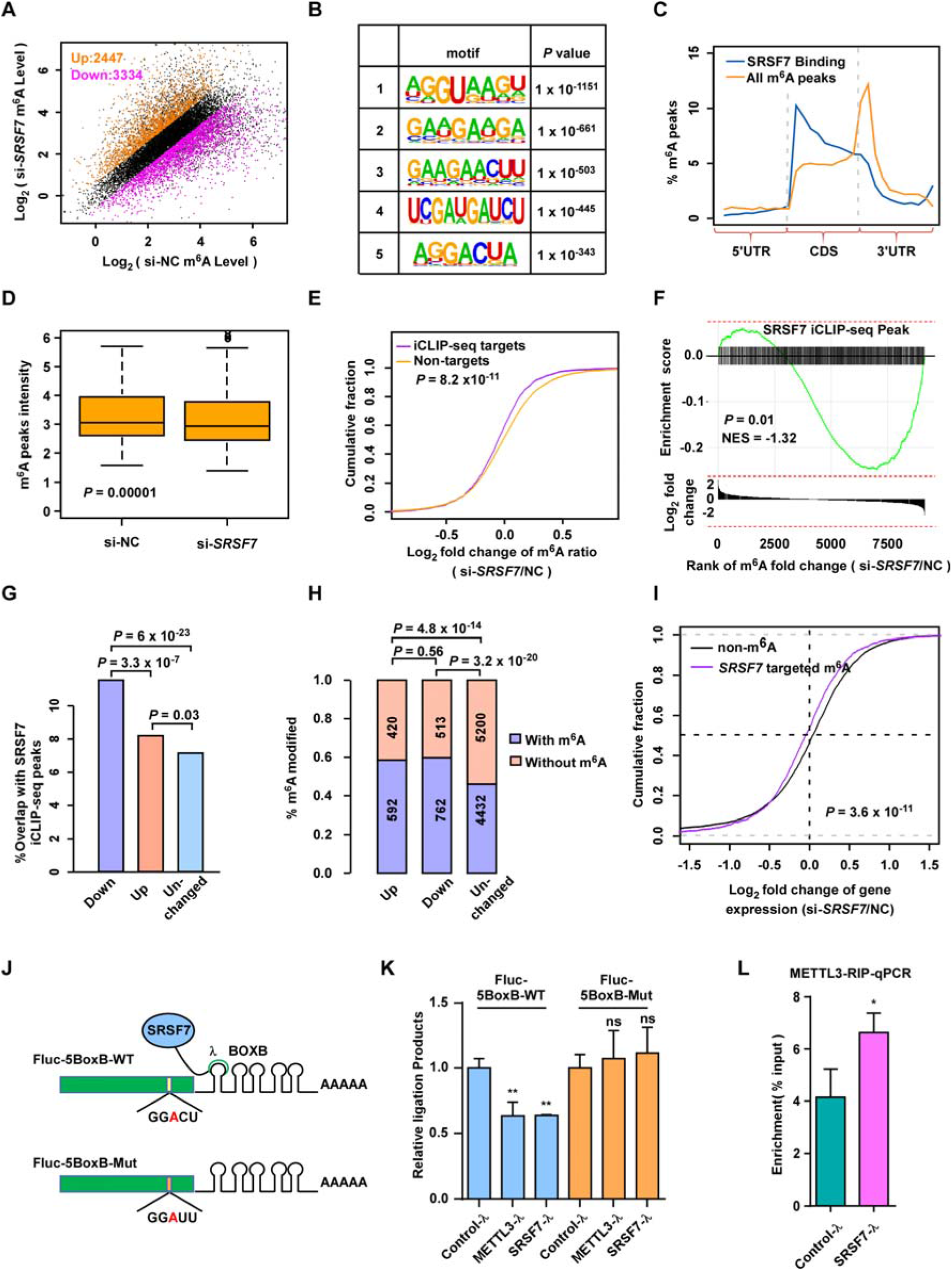
SRSF7 specifically facilitates m^6^A methylation near its binding sites via recruiting METTL3. **A.** Scatter plots showing the up-regulated (orange) and down-regulated (purple) m^6^A peaks in si-*SRSF7* as compared with si-NC in U87MG cells. The numbers of the up-regulated and down-regulated peaks are indicated. **B.** The most significantly enriched motifs in the iCLIP-seq identified SRSF7 binding peaks. **C.** Normalized distributions of m^6^A peaks and SRSF7 iCLIP-seq peaks across 5’UTR, CDS, and 3’UTR of mRNA. **D.** Box plot comparing the m^6^A ratios of the SRSF7 targeted m^6^A peaks in control and SRSF7-KD U87MG cells. **E.** Plot of cumulative fraction of log2 fold change of m^6^A ratios upon *SRSF7* knockdown using si-*SRSF7* for the m^6^A peaks overlap or non-overlap with SRSF7 iCLIP-seq peaks. *P* value of two-tailed Wilcoxon test is indicated. **F.** Plot of GSEA analysis displaying the distribution of SRSF7 iCLIP-seq peaks (upper panel) across the m^6^A peaks ranked by log2 fold change of m^6^A ratios upon *SRSF7* knockdown (si-*SRSF7*) (lower panel). The m^6^A peaks overlap SRSF7 iCLIP-seq peaks are indicated by vertical line in the upper panel. The *P* value and normalized enrichment score (NES) of GSEA are indicated. **G.** Bar plot comparing the percentages of m^6^A peaks overlapped with SRSF7 iCLIP-seq peaks for down-regulated, up-regulated, and unchanged m^6^A peaks upon *SRSF7* knockdown respectively. The pairwise *P* values of two-tailed Chi-square tests are indicated at the top. **H.** Bar plot comparing the percentages of m^6^A modified genes for genes with down-regulated, up-regulated, and unchanged gene expression upon *SRSF7* knockdown respectively. The pairwise *P* values of two-tailed Chi-square tests are indicated at the top. **I.** Plot of cumulative fraction of log2 fold change of gene expression upon *SRSF7* knockdown for unmethylated genes and genes with SRSF7 targeted m^6^A peaks respectively. *P* value of two-tailed Wilcoxon test is indicated. **J.** Schematic diagram displaying the construct of the SRSF7 tethering assay with GGACU m^6^A motif (upper) and disruptive GGAUU motif (lower). **K.** Bar plot comparing the SELECT method measured relative ligation product, which anti-correlated with the m^6^A level, for the m^6^A site in F-luc-5BoxB without or with mutation in the m^6^A motif in U87MG cells transfected with Control-λ, SRSF7-λ, and METTL3-λ respectively. Data are presented as mean ± SEM, n=3. ** *P* <0.01, ns, no significant difference. One-way ANOVA with Dunnett’s post hoc test. **L.** Bar plot comparing the METTL3 RIP-qPCR enrichment of the F-luc mRNA in U87MG cells transfected with SRSF7-λ and Control-λ respectively. Data are presented as mean ± SEM, n=3. * *P* <0.05. Student’s two-tailed *t* test.

To further confirm that SRSF7 regulates the m^6^A sites through binding near the m^6^A sites, we performed iCLIP-seq [37] for SRSF7 to identify the transcriptome-wide binding sites of SRSF7 in U87MG cells. We identified 40476 iCLIP-seq peaks using CTK took kit [38] (Table S1). The enriched motifs were similar as the previously reported motif of SRSF7 (GAYGAY) [39], suggesting the high reliability of our iCLIP-seq data (Figure 2B).

Interestingly, the m^6^A motif were also enriched in the SRSF7 iCLIP-seq peaks (Figure 2B), suggesting the co-localization of SRSF7 with m^6^A sites. We found only 7.9% and 3.1% of the peaks were in introns and noncoding RNAs respectively; in contrast, 66.9% of the peaks were in protein-coding regions, which are similar as the distribution of m^6^A (Figure S2G). However, the peaks were more enriched at the 5’ end of the protein-coding regions, which was distinct from m^6^A peaks; while the peaks colocalized with m^6^A peaks were enriched at both 5’ end and 3’end, further suggesting that SRSF7 specifically regulates only a portion of m^6^A peaks other than global regulation (Figure 2C and Figure S2H).

We were then interested in whether SRSF7 binding were related to the m^6^A alteration due to SRSF7 depletion. We found that although the overall m^6^A ratios of all m^6^A peaks do not change upon SRSF7 knockdown, the m^6^A ratios of m^6^A peaks colocalized with SRSF7 siCLIP-seq peaks were significantly down-regulated upon SRSF7 knockdown, suggesting SRSF7 can only promote the m^6^A near its binding sites (Figure 2D). As compared with the m^6^A peaks unbound by SRSF7, the m^6^A ratio of SRSF7 bound m^6^A peaks were significantly down-regulated due to *SRSF7* knockdown, indicating that SRSF7 specifically facilitates the m^6^A near its binding sites (Figure 2E). As shown in Figure 2F, we also revealed significant enrichment of SRSF7 iCLIP-seq peaks in (or overlap with) the down-regulated m^6^A peaks upon *SRSF7* knockdown. In addition, the SRSF7 binding sites were significantly enriched in m^6^A peaks down-regulated upon *SRSF7* knockdown as compared with the up-regulated and unchanged m^6^A peaks, further supporting that SRSF7 binding results in locally enhanced other than decreased m^6^A methylation (Figure 2G). On the other hand, although the module was constructed from diverse cell lines, the SRSF7 binding sites in U87MG cells were still marginally significantly enriched (*P* = 0.03) in the orange module, which is a larger module merged by M5 and other 4 correlated modules, as compared with other modules. The m^6^A peaks in the orange module were also significantly down-regulated upon *SRSF7* knockdown as compared with the m^6^A peaks in other modules, suggesting SRSF7 promotes the m^6^A of this module (Figure S2I).

### SRSF7 significantly regulates gene expression through regulating m^6^A

We then studied whether SRSF7 affects the gene expression through regulating m^6^A in U87MG cell. The expression of 1012 and 1275 genes were up-regulated and down-regulated respectively due to *SRSF7* knockdown (Figure S3A). GO enrichment analyses found that the down-regulated genes were enriched in terms such as cell division, cell migration, cell cycle, consistent with the GO terms enriched in differentially methylated genes (Figure S3B).

However, the up-regulated genes were enriched in terms macroautophagy, vesicle docking, protein transport, which were quite different from the GO terms enriched in differentially methylated genes (Figure S3C). Gene set enrichment analysis (GSEA) also support the gene expression changes were involved in cell division, cell cytoskeleton and cell cycle (Figure S3D-F). We found both the up-regulated genes and down-regulated genes significantly enriched for m^6^A modified genes as compared with the genes without expression change (*P* = 4.8 × 10^-14^ for up-regulated genes; *P* = 3.2 × 10^-20^ for down-regulated genes; two-tailed Chi-square test; Figure 2H). This result suggests SRSF7 can both up-regulate and down-regulate gene expression through m^6^A, consistent with the previous reports that m^6^A has dual effects on gene expression depends on how these m^6^A sites are recognized by diverse m^6^A readers [23, 27–29]. To further clarify the direct effects of SRSF7, we investigated the effects of SRSF7 binding on gene expression though regulating m^6^A. As shown in Figure 2I, the genes with SRSF7 targeted m^6^A peaks are overall significantly down-regulated as compared with non-modified genes upon SRSF7 knockdown (*P* = 3.6 × 10^-11^, two-tailed Wilcoxon test, Figure 2I).

### Artificially tethering SRSF7 on RNA directs *de novo* m6A methylation through recruiting METTL3

We then performed a tethering assay to test whether direct tethering of SRSF7 protein was sufficient to dictate the m^6^A modification nearby in U87MG cells. For this purpose, we respectively fused the full-length CDS region SRSF7 and METTL3 with λ peptide, which can specifically recognize BOX B RNA [40]. We utilized a previously established F-luc-5BoxB luciferase reporter, which has five Box B sequence in the 3’UTR and a m^6^A motif (GGACU) 73 bp upstream of the stop codon (Figure 2J) [30]. We found tethering SRSF7 and METTL3 could both significantly up-regulate the modification of m^6^A site on the reporter to the similar degree using SELECT method [41], indicating that SRSF7 can similarly dictate the methylation of nearby m^6^A site as METTL3 (Figure 2K). A disruptive synonymous point mutation in the m^6^A motif, which changes the GGA<u>C</u>U to GG<u>A</u>UU, completely disrupted the effects of m^6^A change by tethering SRSF7 and METTL3 respectively, indicating the high reliability of the tethering assay (Figure 2K). In addition, we found that binding of METTL3 on F-luc RNA was significantly up-regulated when tethered SRSF7 to F-luc-5BoxB, indicating that SRSF7 promotes the installation of m^6^A through recruiting METTL3 (Figure 2L).

### SRSF7 specifically targets and facilitates the methylation of m^6^A on genes involved in cell proliferation and migration

Since SRSF7 iCLIP-seq peaks are significantly enriched in down-regulated m^6^A peaks upon SRSF7 knockdown (Figure 2G), to further dissect the specific m^6^A targets that directly regulated by SRSF7 binding, we intersected the 40476 SRSF7 iCLIP-seq peaks and 3334 down-regulated m^6^A peaks upon SRSF7 knockdown to obtained 911 SRSF7 directly regulated m^6^A peaks in 760 genes (Figure 3A and Table S2). As shown in Figure 3B, the distribution of SRSF7 directly regulated m^6^A peaks was still similar as the canonical distribution of m^6^A peaks, suggesting SRSF7 are not accounting for the formation of the canonical topology of m^6^A like VIRMA [30]. Gene Otology analysis revealed that the genes with SRSF7 directly regulated m^6^A peaks were mainly involved in cell migration, cell adhesion, cell proliferation, glioma, cell cycle and pathway in cancer (Figure 3C and Figure S4A). In contrast, the genes with SRSF7 iCLIP-seq peaks not colocalize with m^6^A peaks were enriched in totally different terms which were not directly related to cell proliferation and migration (Figure S4B). The results suggest that the elevated expression of SRSF7 in GBM patients may involve in migration and proliferation of the cancer cells through regulating the m^6^A of corresponding genes.

**Figure 3.**
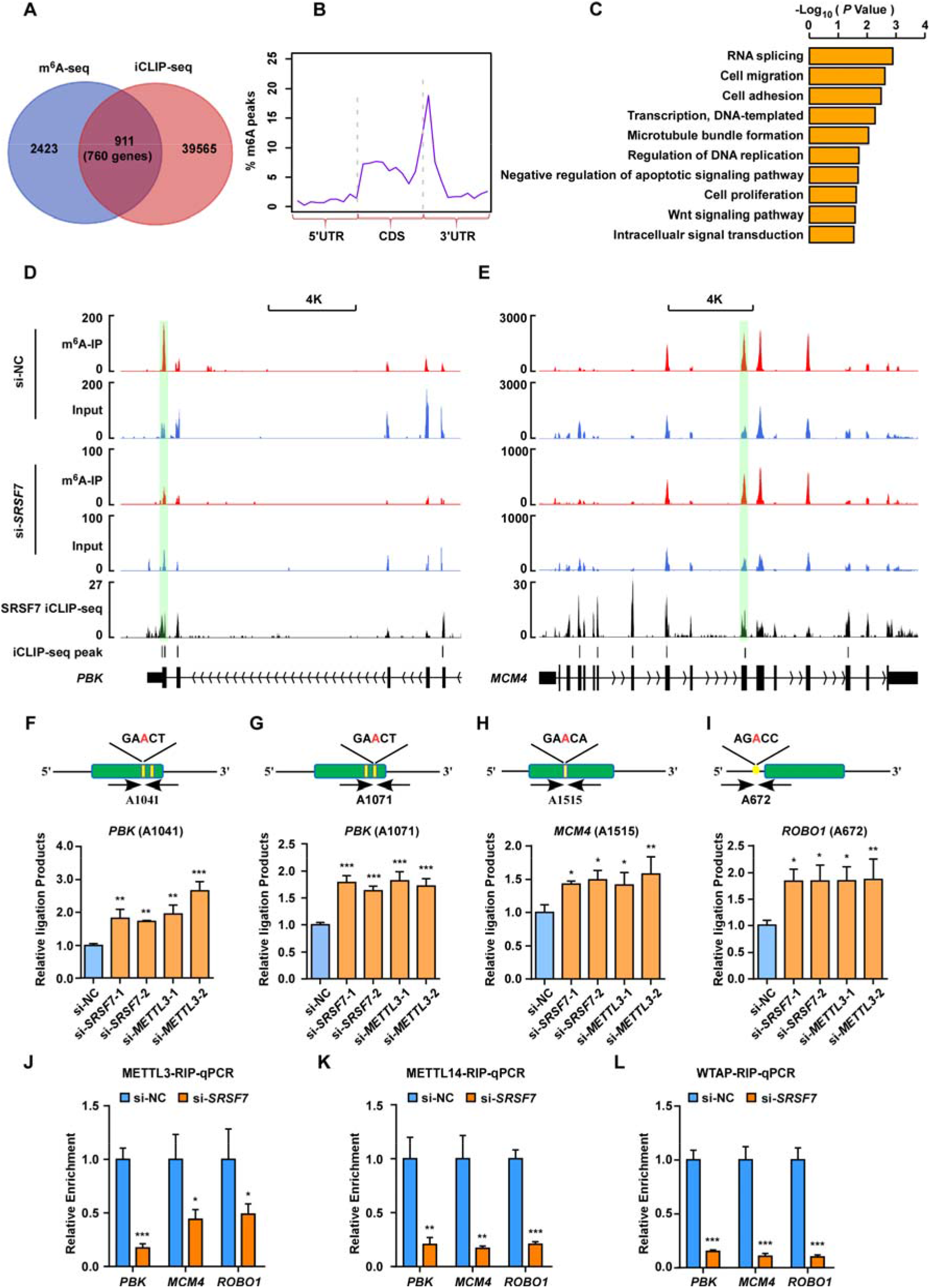
SRSF7 specifically targets and facilitates the methylation of m^6^A on genes involved in cell proliferation and migration. **A.** Venn diagram showing the overlapping of down-regulated m^6^A peaks upon *SRSF7* knockdown and SRSF7 iCLIP-seq peaks. **B.** Normalized distribution of the overlapped m^6^A peaks in (a) across 5’UTR, CDS, and 3’UTR of mRNA. **C.** GO enrichment of the corresponding genes with the overlapped m^6^A peaks in (A). **D, E.** Tracks displaying the read coverage of IPs and inputs of m^6^A-seq as well as the SRSF7 iCLIP-seq on *PBK*, *MCM4*. The SRSF7 directly regulated m^6^A peaks are highlighted. The y-axes of NC and si-SRSF7 were differently used to intuitionally indicate the m^6^A differences other than expression differences. **F-I.** Validation of m^6^A changes using SELECT method of single-nucleotide m^6^A sites on *PBK* at 1041 and 1071 (F-G), *MCM4* at 1515 (H), *ROBO1* at 672 (I) in U87MG cells transfected with scramble (si-NC) and 2 siRNAs of SRSF7 (si-*SRSF7*-1, si-*SRSF*-2), and 2 siRNAs of METTL3 (si-*METTL3*-1, si-*METTL3*-2) respectively. The tested m^6^A motifs are indicated on the schematic structures of mRNAs at the top panels. The green boxes represent protein-coding regions, the thin lines flanking the green boxes represent UTR regions. Arrows indicate the primers for SELECT. Data are presented as mean ± SEM, n=3. * *P* <0.05, ** *P* <0.01, *** *P* <0.001. One-way ANOVA with Dunnett’s post hoc test. **J-L.** Bar plot comparing the RIP-qPCR determined relative enrichment of METTL3 (J), METTL14 (K), and WTAP (L) binding to the mRNA of *PBK*, *MCM4*, and *ROBO1* in control and *SRSF7*-KD U87MG cells. Data are presented as mean ± SEM, n=3. * *P* <0.05, ** *P* <0.01, *** *P* <0.001. Student’s two-tailed *t* test.

To further validate the 911 SRSF7 directly regulated m^6^A peaks, we then selected 3 m^6^A peaks in 3 tumorigenic genes involved in migration or proliferation of GBM. All of the 3 peaks on *PBK*, *MCM4*, and *ROBO1* were successfully validated (the signal tracks of these m^6^A peaks were demonstrated in Figure 3D-E and Figure S4C). We detected 4 single-nucleotide m^6^A sites in the 3 m^6^A peaks according to the public available miCLIP-seq data [42, 43]. The methylation levels of the 4 m^6^A sites in the 3 m^6^A peaks (*PBK* at 1041 and 1071, *MCM4* at 1515, *ROBO1* at 672) were significantly decreased upon *SRSF7* knockdown and *METTL3* knockdown respectively based on SELECT method [41], indicating SRSF7 has similar effects of promoting m^6^A as METTL3 on these selected m^6^A sites (Figure 3F-I). We also found that the binding efficiencies of METTL3, METTL14, and WTAP on the RNAs of these 3 genes were significantly reduced upon *SRSF7* knockdown based on RIP-qPCR (Figure 3J-L). Collectively, these results show that SRSF7 promotes m^6^A modification on tumorigenic genes through recruiting METTL3.

### SRSF7 promotes proliferation and migration of glioblastoma cells partially dependent on METTL3

Since SRSF7 specifically regulates the m^6^A of tumorigenic genes in GBM cells, we therefore wanted to confirm whether it plays important roles in GBM. We found the expression of *SRSF7* was highly elevated in glioma specimens, especially in glioblastoma (grade IV) tissues according to CGGA data (Figure 4A), which was confirmed by Immunohistochemistry (IHC) in human glioma tissues (Figure 4B) and consistent with previous report [10]. To further confirm this finding, we tested the expression of SRSF7 in 11 GBM cell lines as well as normal human astrocytes (NHAs). We found the mRNA expression of SRSF7 was significantly elevated in most of the glioma cell lines and the protein level was highly expressed in all glioma cell lines as compared with NHA (Figure 4C and D).

**Figure 4.**
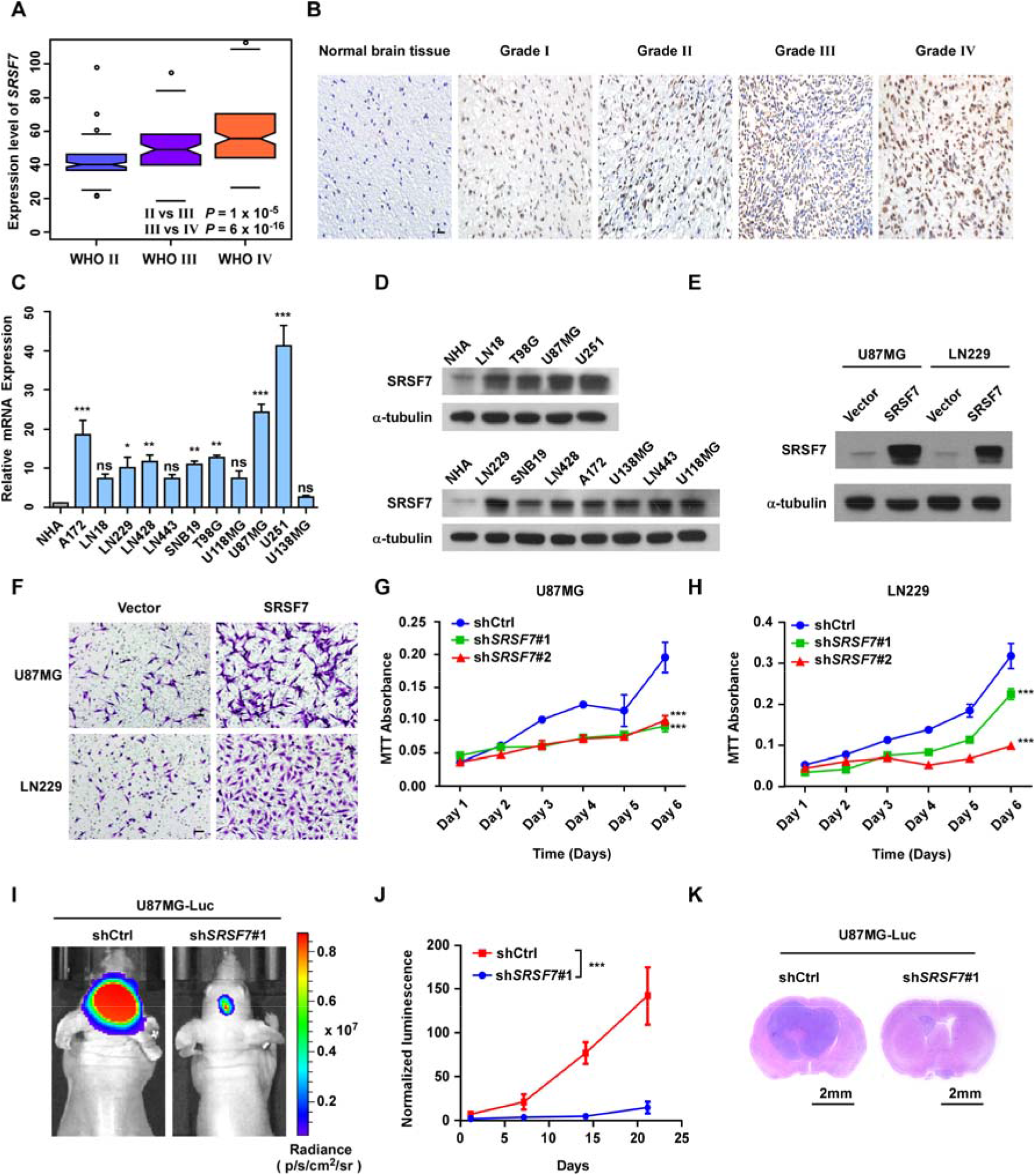
SRSF7 promotes proliferation and migration of glioblastoma cells. **A.** Boxplot comparing the expression of *SRSF7* during GBM patients of different stages from CGGA dataset. *P* values of two-tailed Student’s t test are indicated. **B.** Left: IHC staining of SRSF7 in normal brain and glioma specimens, Scale Bar: 20 μM. **C.** Bar plot comparing the mRNA expression levels of *SRSF7* in 11 GBM cell lines and Normal Human Astrocytes (NHA). Data are presented as mean ± SEM (standard error of mean), n=2. * *P* < 0.05, ** *P* < 0.01, *** *P* < 0.001, ns, no significant difference. One-way ANOVA with Dunnett’s post hoc test. **D.** Western blot comparing the protein levels of SRSF7 in 11 GBM cell lines and NHA. **E.** Western blot showing efficiently overexpression of *SRSF7* in U87MG and LN229 cells. **F.** Representative images of transwell migration assay in U87MG and LN229 cells overexpressing *SRSF7*. Scar bars: 50 μm. **G, H.** The cell viability of *SRSF7* knockdown and control in U87MG (G) and LN229 (H) cells were measured by MTT assay at indicated time point. Data are presented as mean ± SEM, n = 3. *** *P* < 0.001. Two-way ANOVA with Dunnett’s post hoc test. **I.** Representative bioluminescence images of mice-bearing the intracranial glioma xenografts formed by U87MG cells transduced with control shRNA or *SRSF7* shRNA respectively. **J.** Line graph showing the normalized luminescence of intracranial glioma xenografts tumors formed by U87MG cells transduced with control shRNA or *SRSF7* shRNA respectively. Data are presented as mean ± SEM, n = 6. *** *P* < 0.001. Student’s two-tailed *t* test. **K.** Representative images of H&E staining of glioma tissue sections from indicated mice, Scale Bar: 2mm.

Because the genes with SRSF7 directly regulated m^6^A peaks were enriched in cell proliferation and migration related GO terms (Figure 3C), we knocked down *SRSF7* in U87MG cells and LN229 cells and performed EdU, colony formation, and transwell assays to test the effects of SRSF7 on cell proliferation and migration. We found overexpression of *SRSF7* prompted the cell proliferation and migration of these two cell lines (Figure 4E-F and Figure S5A). Consistently, depletion of *SRSF7* significantly impaired the proliferation and migration in U87MG and LN229 cell lines (Figure 4G-H and Figure S5B-D) and overexpression of *SRSF7* can rescue the inhibition of proliferation and migration caused by *SRSF7* knockdown (Figure S5E-G), which were similar as the effects of *METTL3* knockdown in the same cell lines [44, 45]. Although METTL3 has been reported to regulate the stemness of GBM cells [44–47], the genes with SRSF7 directly regulated m^6^A peaks have no enrichment of stemness related terms (Figure 3C). Here, we found neither knockdown or overexpression of *SRSF7* could affect the neurosphere formation in U87MG cells, which suggesting that SRSF7 plays more specific roles in GBM than METTL3 through specific regulation of m^6^A (Figure S5H). To investigate the oncogenic role of SRSF7 in GBM cells *in vivo*, we utilized an intracranial xenograft tumor model, in which we transplanted *SRSF7* depleted as well as control U87MG stable cell lines into the nude mice. Consistent with the *in vitro* findings, *SRSF7* knockdown significantly inhibited the growth of glioma xenografts (Figure 4I-K). We further confirmed that SRSF7 cannot regulate the gene or protein expression of the core methyltransferase complex (Figure 5A and Figure S6A-E), and METTL3 or WTAP cannot regulate the expression of SRSF7 either in U87MG or LN229 cells (Figure S6F-G). In addition, SRSF7 knockdown did not change the nuclear speckle localization of METTL3, METTL14, or WTAP (Figure S6H-J). The above results indicate that SRSF7 promotes the proliferation and migration, which are usually related to oncogenic roles, of GBM cells.

**Figure 5.**
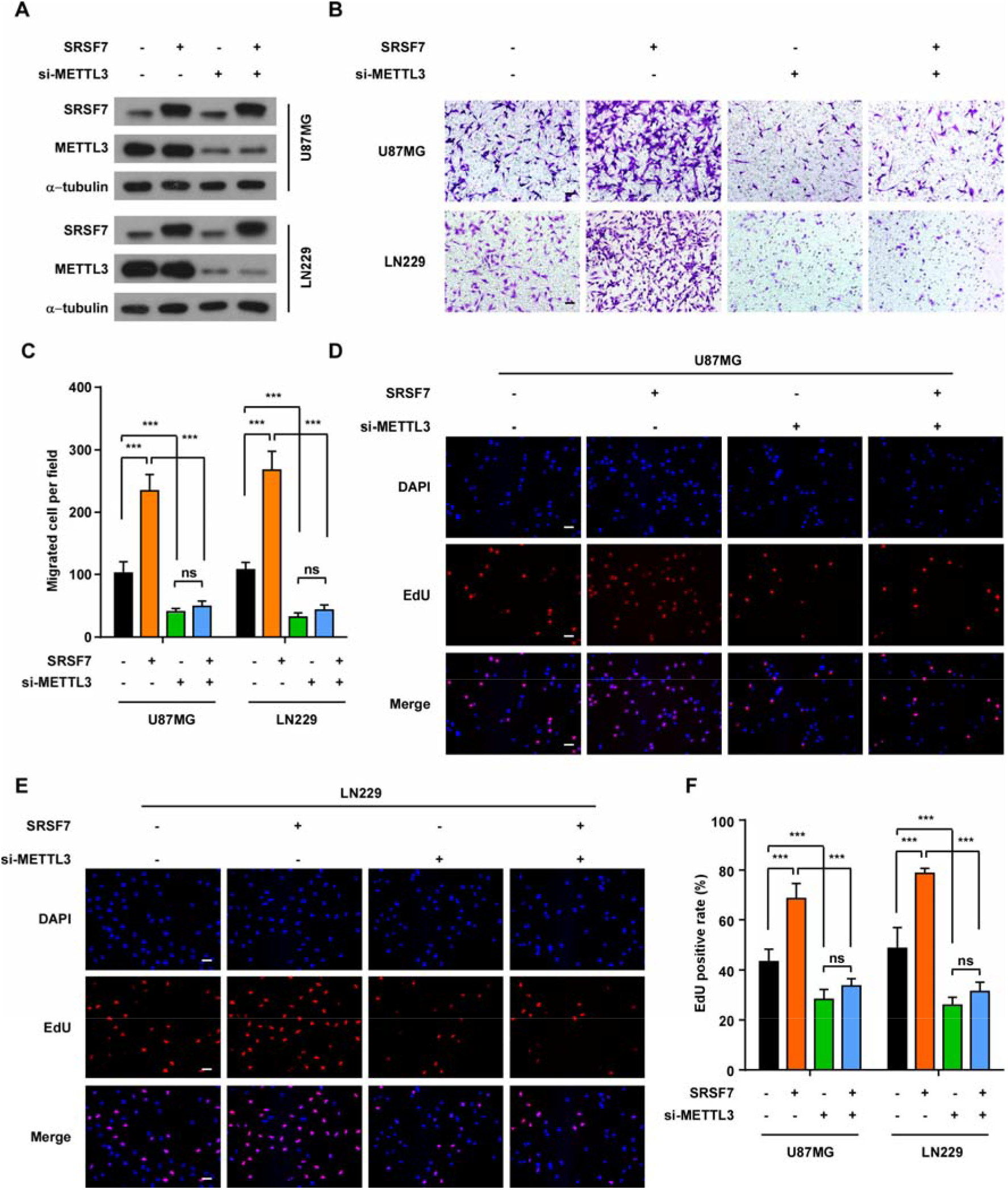
SRSF7 promotes the proliferation and migration of glioblastoma cells partially dependent on METTL3. **A.** Western blot showing the protein level of SRSF7 and METTL3 in SRSF7 overexpressed U87MG and LN229 cells transfected without or with si-*METTL3*-1 cell as indicated. **B, C.** Representative images (B) and bar plot (C) comparing the number of migrated cells in transwell migration assay in SRSF7 overexpressed U87MG and LN229 cells transfected without or with si-*METTL3*-1 cell as indicated. Data are presented as mean ± SEM, n = 5. *** *P* < 0.001. ns, no significant difference. One-way ANOVA with Tukey’s post hoc test. Scar bars: 50 μm. **D-E.** Representative images of EdU staining in SRSF7 overexpressed U87MG (D) and LN229 (E) cells transfected without or with si-*METTL3*-1 as indicated. Scar bars: 50 μm. **F.** Bar plot comparing the EdU positive rate of EdU staining in SRSF7 overexpressed U87MG and LN229 cells transfected without or with si-*METTL3*-1 as indicated. Data are presented as mean ± SEM, n = 5. * *P* < 0.05, *** *P* < 0.001. ns, no significant difference. One-way ANOVA with Tukey’s post hoc test.

It was reported that METTL3 plays oncogenic roles in GBM [48–51], we were therefore interested in whether SRSF7 plays oncogenic roles through specifically guiding METTL3 to oncogenic genes. We found *METTL3* knockdown largely, although not completely, disrupted the effects of *SRSF7* overexpression on promoting the migration (Figure 5A-C) and proliferation (Figure 5D-F) of U87MG and LN229 cell, indicating that SRSF7 regulates migration and proliferation partially depends on METTL3. The above results are consistent with our model that SRSF7 specifically guides METTL3 to the specific oncogenes and METTL3 takes in charge to install the m^6^A on these RNAs.

### SRSF7 promotes the proliferation and migration of GBM cells partially through the m^6^A on *PBK* mRNA

We were then interested in the downstream targets of SRSF7 that mediated the proliferation and migration changes of GBM cells *via* m^6^A. Out of the 760 genes with SRSF7 directly regulated m^6^A peaks, *PBK* is the most significantly down-regulated gene upon *SRSF7* knockdown. Meanwhile, as shown in Figure 3D and Figure 3F-G, we have confirmed that SRSF7 knockdown significantly reduced the m^6^A of two m^6^A sites on PBK (A1041 and A1071). PBK is also a serine/threonine protein kinase which is aberrantly overexpressed in various cancers and plays important roles in promoting the proliferation and migration of multiple cancers including glioma [52–56]. Based on the CGGA dataset, *PBK* is significantly higher expressed in WHO IV of glioma patients as compared with WHO II and WHO III, the highly expression of PBK is significantly associated with poor prognosis in GBM (Figure S7A-B). Furthermore, the gene expression of *PBK* is positively correlated with *SRSF7* and *METTL3* based on CGGA dataset, suggesting a regulatory role between them (**Figure 6**A and Figure S7C). We found that overexpression of PBK could partially rescue the *SRSF7* knockdown induced inhibition of proliferation and migration of U87MG and LN229 cells, indicating that PBK is an important downstream target of SRSF7 and partially mediated the effects of SRSF7 on promoting the proliferation and migration of GBM cells (Figure 6B-C and Figure S7D-E). We were therefore interested in whether and how the expression of *PBK* was regulated by SRSF7.

**Figure 6.**
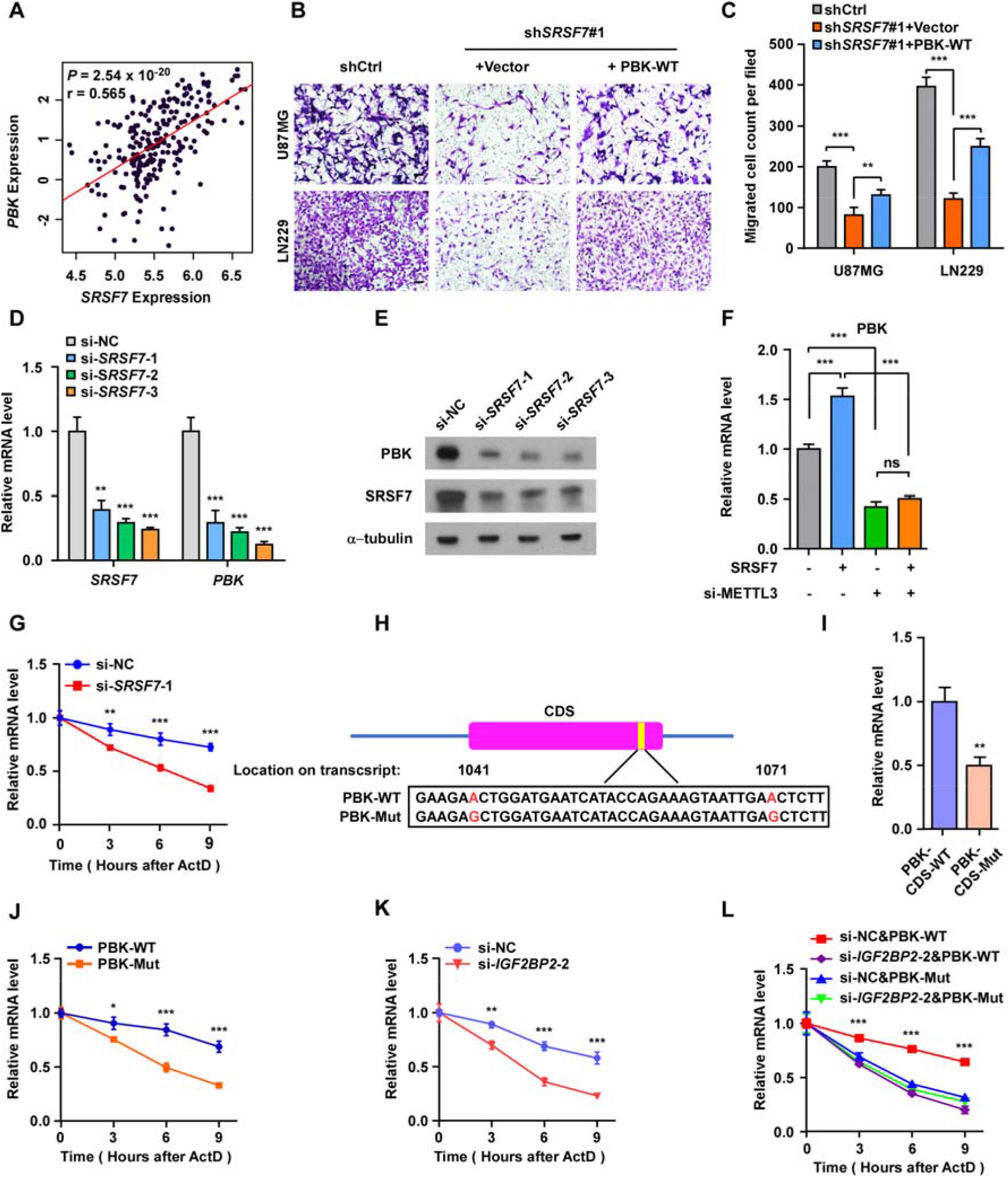
SRSF7 promotes the proliferation and migration of GBM cells partially through the m^6^A on *PBK* mRNA. **A.** Scatter plot showing the correlation between *SRSF7* and *PBK* gene expression across GBM patients from CGGA dataset, the *P* value and correlation coefficient are indicated. **B, C.** Representative images (B) and bar plot (C) comparing the number of migrated cells in transwell migration assay in U87MG and LN229 cells upon *SRSF7* knockdown and rescue by co-transducing full-length WT *PBK* CDS regions. Data are presented as mean ± SEM, n = 5. ** *P* < 0.01, *** *P* < 0.001. One-way ANOVA with Tukey’s post hoc test. Scar bars: 50 μm. **D.** Bar plot showing the relative mRNA level of *SRSF7* and *PBK* in U87MG cells transfected with scramble (si-NC) and 3 different siRNAs of *SRSF7* respectively. Data are presented as mean ± SEM, n = 3. ** *P* < 0.01, *** *P* < 0.001. Student’s two-tailed *t* test. **E.** Western bolt comparing the protein levels of SRSF7 and PBK in U87MG cells transfected with scramble (si-NC) and 3 different siRNAs of *SRSF7* respectively. **F.** Bar plot showing the relative mRNA level of *PBK* in SRSF7 overexpressed U87MG cells transfected without or with si-*METTL3*-1 as indicated. Data are presented as mean ± SEM, n = 3. *** *P* < 0.001, ns, no significant difference. One-way ANOVA with Tukey’s post hoc test. **G.** Relative mRNA levels of *PBK* after actinomycin D treatment at indicated time points in U87MG cells transfected with scramble (si-NC) and siRNA of *SRSF7* respectively. Data are presented as mean ± SEM, n = 3. ** *P* < 0.01, *** *P* < 0.001. Two-way ANOVA with Bonferroni’s post hoc test. **H.** Schematic diagram of mutation of the two m^6^A site in the *PBK* CDS region. **I.** Relative mRNA level of *PBK* in U87MG cells transfected with full-length WT or mutant (Mut) *PBK* CDS regions for 48 hours. Data are presented as mean ± SEM, n = 3. ** *P* < 0.01. Student’s two-tailed *t* test. **J.** Relative mRNA levels of *PBK* after actinomycin D treatment at indicated time points in U87MG cells transfected with full-length WT or mutant (Mut) *PBK* CDS regions respectively. Data are presented as mean ± SEM, n = 3. ** *P* < 0.01, *** *P* < 0.001 Two-way ANOVA with Bonferroni’s post hoc test. **K.** Relative mRNA levels of *PBK* after actinomycin D treatment at indicated time points in U87MG cells transfected with scramble (si-NC) and siRNA of *IGF2BP2* (si-*IGF2BP2*) respectively. Data are presented as mean ± SEM, n = 3. ** *P* < 0.01, *** *P* < 0.001. Two-way ANOVA with Bonferroni’s post hoc test. **L.** Relative mRNA levels of *PBK* after actinomycin D treatment at indicated time points in WT *PBK* or Mut *PBK* overexpressed U87MG cells transfected with scramble (si-NC) and siRNA of *IGF2BP2* respectively. Data are presented as mean ± SEM, n = 3. *** *P* < 0.001. Two-way ANOVA with Dunnett’s post hoc test.

First, we tested whether SRSF7 played a regulator role on *PBK* through regulating its m^6^A. We found that *SRSF7* knockdown significantly decreased the mRNA and protein expression of *PBK* in U87MG cells (Figure 6D and E). Overexpression of *SRSF7* significantly up-regulated the gene expression of *PBK*, and *METTL3* knockdown largely disrupted the effect of SRSF7 on the expression of *PBK* in U87MG cells, indicating that SRSF7 regulates *PBK* depends on METTL3 (Figure 6F and Figure S7F-G).

We then asked how the m^6^A of *PBK* affects its expression. We found *SRSF7* knockdown also significantly promoted the degradation of *PBK* mRNAs, suggesting that SRSF7 increase *PBK* gene expression through promoting the stability of *PBK* mRNAs (Figure 6G). To further confirm this regulation of RNA stability depends on the m^6^A of *PBK*, we introduced two synonymous A to G mutations to disrupt the two m^6^A sites on *PBK* (Figure 6H). We found the overexpression of *PBK* mutants exhibited significantly lower expression and lower stability of *PBK* mRNA than overexpression of wild type *PBK*, suggesting the modification of the two m^6^A sites on *PBK* are essential for the stability of *PBK* mRNA (Figure 6I and J).

Because m^6^A readers IGF2BP1-3 have been reported to promote the stabilities of mRNAs and play oncogenic roles in multiple cancers [27]. We then tested whether IGF2BP2, a gene significantly up-regulated in GBM, could affect the RNA stability of *PBK* through binding the m^6^A sites. We found knockdown of *IGF2BP2* decreased the expression and stability of endogenous *PBK* mRNA (Figure 6K and Figure S7H), which is consistent with the finding that the gene expression of *IGF2BP2* is positively correlated with *PBK* based on CGGA dataset (Figure S7I). Knockdown of *IGF2BP2* could also significantly decrease the stability of the exogenous overexpressed wild type PBK other than *PBK* mutants with the two m^6^A sites disrupted, suggesting that the regulatory roles of IGF2BP2 on the stability of *PBK* depends on the two m^6^A sites (Figure 6L).

### SRSF7 regulates m^6^A independent of alternative splicing and polyadenylation

Since SRSF7 was previously recognized as a splicing factor [2–4], to test whether SRSF7 can regulate alternative splicing in U87MG cells, we analyzed the differential alternative splicing of input RNAs between SRSF7 knockdown and control using rMATS [57]. We found 1344 differentially spliced events, including 734 skipped exons (SE), 222 retained introns (RI), 129 alternative spliced 5’ splice sites (A5SS), 173 alternative spliced 3’ splice sites (A3SS), and 86 mutually exclusive exons (MXE). Of note, none of *PBK*, *MCM4*, or *ROBO1* has alternative splicing change upon SRSF7 knockdown. We then used rMAPS2 [58] to study the enrichment of SRSF7 iCLIP-seq peaks near the splice sites of differentially spliced SE events, which are the most abundant type for reliable analyses. We found the iCLIP-seq targets of SRSF7 were significantly enriched in the alternative exons of the differentially spliced evens, suggesting SRSF7 binding directs the splicing changes (Figure 7A). GO analysis revealed that the genes with significant splicing changes were also enriched in functional terms “Cell-cell adhesion”, “Cell cycle”, suggesting SRSF7 can also regulate cell proliferation and migration through alternative splicing (Figure 7B). For the 760 genes with

**Figure 7.**
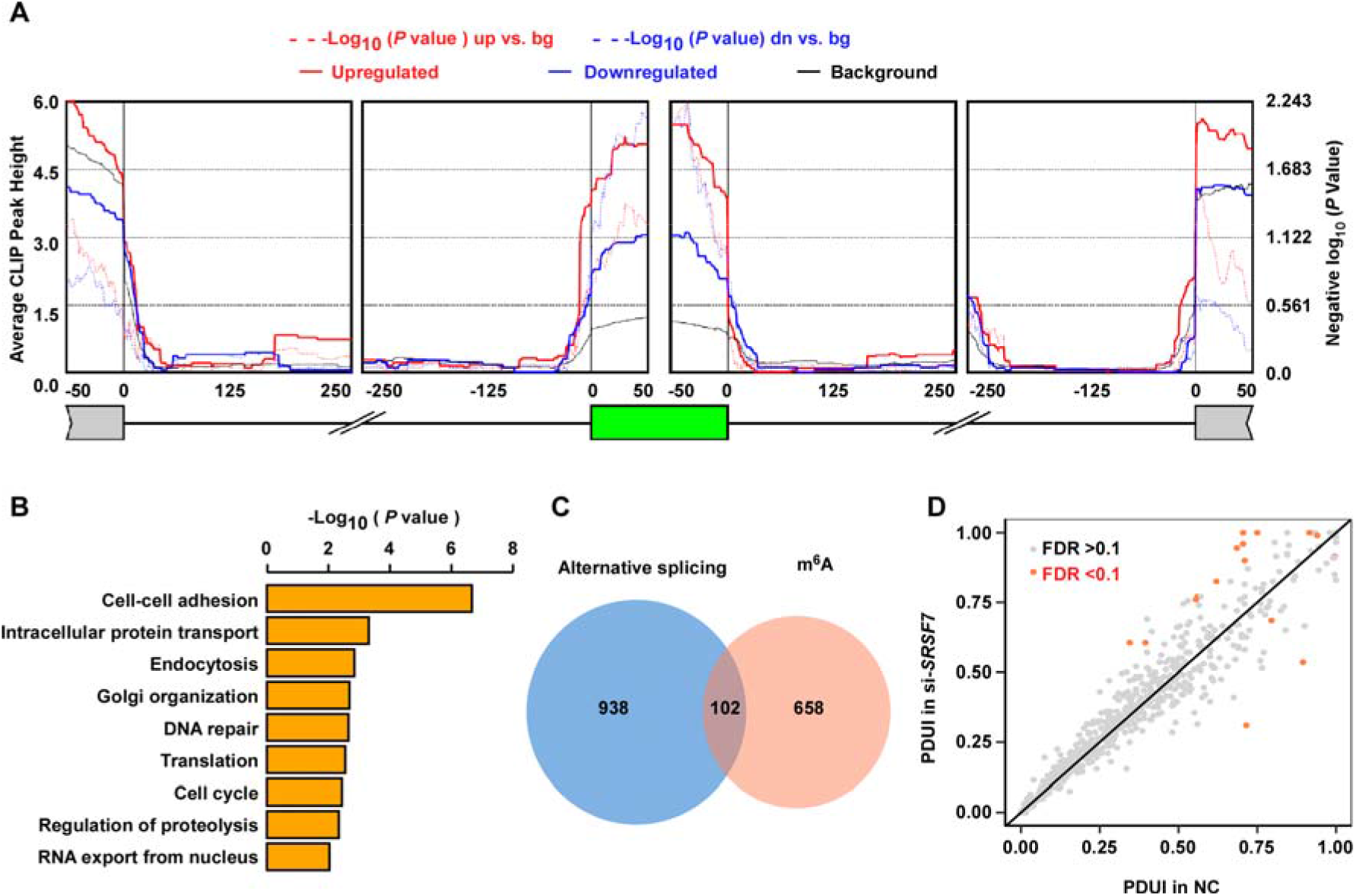
SRSF7 regulates m^6^A independent of alternative splicing and polyadenylation. **A.** rMAPS2 generated metagene plot showing the enrichment of SRSF7 iCLIP-seq peaks at the regions around corresponding splice sites of the differentially spliced SE events upon *SRSF7* knockdown. **B.** GO enrichment of differentially spliced genes (all types) upon *SRSF7* knockdown. **C.** Venn diagram showing the overlap between differentially spliced genes (all types) and genes with SRSF7 directly regulated m^6^A peaks. **D.** Scatter plot comparing the distal poly(A) site usage index (PDUI) between control and SRSF7 knockdown in U87MG cells.

SRSF7 directly regulated m^6^A, only 102 (13.4%) of them had significant splicing changes upon *SRSF7* knockdown (Figure 7C), which represented a non-significant overlap that could easily occur by random chance (P = 0.3, two tailed Chi-square test). For the 129 m^6^A peaks in the 102 genes, only 36 peaks in 28 genes were localized within the local regions of differentially splicing events spanning between upstream exons to downstream exons, of which only 7 m^6^A peaks were located within the alternative exons or regions. The above results indicate that SRSF7 regulates m^6^A and alternative splicing independently through distinct binding sites, consistent with our observation that only a part of SRSF7 proteins co-localize with METTL3 and only a part of SRSF7 binding sites can regulate m^6^A.

Since SRSF7 was also reported to regulate alternative polyadenylation (APA) of RNAs [5], we also analyzed the differential APAs of input RNAs between *SRSF7* knockdown and control in U87MG cells using DaPars [59]. We only found 14 APA events were significantly changed (Figure 7D), and none of the SRSF7 directly regulated m^6^A peaks was located within the 14 APA regions regulated by SRSF7, suggesting noninterference between SRSF7 regulated m^6^A and APA.

## Discussion

m^6^A has been reported to play important roles in diverse systems through different targets, there are widespread m^6^A sites on most of the genes with diverse functions, it is very important for cells to dynamically coordinate the methylation of different genes to fulfil specific functions. In this study, we found SRSF7 specifically regulate the m^6^A on genes involved in cell proliferation and migration, it demonstrated an important role of RBP-mediated specific regulation of m^6^A in co-regulating and coordinating a batch of related m^6^A sites in order to modulate the specific functions in cells. These diverse specific m^6^A regulators provide a versatile toolkit for cells to deal with various inner and outer stimulates. On the other hand, widespread involvement of RBPs in regulating m^6^A suggests that the m^6^A signaling pathways are deeply involved in the regulatory network of genes. Therefore, other signaling or regulatory pathways can modulate the m^6^A through regulating the RBPs in order to fulfil the downstream functions. It is very possible that more and more important functional roles of RBP-mediated specific regulation of m^6^A will be revealed in the future.

SRSF7 is an adaptor of NXF1, which exports mature RNAs out of nucleus, and plays important roles in coupling RNA alternative splicing and polyadenylation to mRNA export [5]. We revealed a novel role for SRSF7 as a regulator of m^6^A methylation via recruiting METTL3. It is very possible that SRSF7 may also couple m^6^A methylation to mRNA export, in this way the specific RNAs must be methylated before export. RBM15, a component of methyltransferase complex, is also an adaptor of NXF1 [60], furthering suggesting that methylation and export could be linked by a series of m^6^A regulators with RNA binding specificities.

Interaction of SRSF7 with the nucleic m^6^A reader YTHDC1 has been reported by different groups [61, 62]. Xiao *et al* found SRSF7 does not mediate the splicing change regulated by YTHDC1 [61]. While Kasowitz *et al* proposed that YTHDC1 regulates alternative polyadenylation through recruiting SRSF7 [62]. The interactions of SRSF7 with both writers and readers of m^6^A suggest that SRSF7 may also work to coordinate the feedback between writing and reading of m^6^A. On the other hand, although the association between m^6^A and alternative polyadenylation has been reported in multiple studies, the mechanism is not clear yet [30, 62–64]. Our finding that SRSF7 specifically regulates m^6^A may provide a novel potential mechanism that link m^6^A and alternative polyadenylation by SRSF7.

We found SRSF7 knockdown did not affect the overall peak intensities of all m^6^A peaks, but the overall peak intensities of SRSF7 targeted m^6^A peaks were significantly down-regulated upon SRSF7 knockdown. The indicated fact that SRSF7 only regulates a small portion of m^6^A sites may be a general feature of all specific regulators of m^6^A, it represents the advantage of using specific m^6^A regulators for cells that require precise regulation of a small portion of m^6^A targets. As we have previously proposed, the specific regulators of m^6^A may work in a similar way as splicing factors [35, 65], which usually do not affect the global splicing levels but a small portion of cell-specific splicing events [34]. On the other hand, although we have proved that only down-regulated m^6^A peaks upon *SRSF7* knockdown enriched for *SRSF7* binding sites, we cannot rule out there are also indirect effects of *SRSF7* knockdown that up-regulates m^6^A, which may counteract the direct effects of SRSF7. We found significant (*P* < 1 × 10^-4^) enrichment of 8 motifs in the up-regulated m^6^A peaks using all m^6^A peaks as background, suggesting that other specific regulators may recruit methyltransferase locally as indirect effects of *SRSF7* knockdown (Figure S8A).

To understand why only a small part of SRSF7 binding peaks can affect m^6^A methylation, we performed motif enrichment analysis for the SRSF7 iCLIP-seq peaks that overlapped with the 911 SRSF7 directly regulated m^6^A peaks using all SRSF7 iCLIP-seq peaks as background. As shown in Supplementary Figure S8B, there are 10 motifs significantly (*P* < 1 × 10^-4^) enriched in the SRSF7 iCLIP-seq peaks that affect m^6^A. The most significantly enriched motifs are m^6^A motifs, suggesting that the existence of m^6^A motif near SRSF7 binding sites are necessary for SRSF7 to promote the m^6^A methylation. This is consistent with our finding that tethering SRSF7 promotes the m^6^A methylation of a nearby m^6^A motif but not the disruptive m^6^A motif with mutation right beside the m^6^A site (Figure 2K). The enrichments of non-m^6^A motifs suggesting that the regulatory role of SRSF7 on m^6^A may be modulated by other factors colocalized with SRSF7 (Supplementary Figure S8B). On the other hand, it was reported that protein modifications of SRSF7 are important for SRSF7 to play different roles on RNA metabolisms. For example, phosphorylated SRSF7 affects RNA splicing, while dephosphorylated SRSF7 promotes nuclear exportation of RNAs [3]. In this study, we found SRSF7 regulated alternative splicing events and alternative polyadenylation events occur independently with m^6^A peaks (Figure 7A-D), suggesting that there is also a comparable fraction of SRSF7 binding sites required for proper alternative splicing and alternative polyadenylation other than m^6^A in GBM cells, probably more sites take charge for nuclear export of RNAs. In addition, not all RBP binding sites reported by CLIP-seq are functional because the binding may not be strong enough. Considering that there are also a small portion of SRSF7 binding sites that can affect alternative splicing, the number of m^6^A regulating SRSF7 binding sites look reasonable for specific regulators that do not affects the nuclear speckle localization of methyltransferase (Figure S6H-I).

m^6^A has been reported to play important roles in cancer development [48–51]. Global disruption of m^6^A by METTL3 depletion has been found to affect tumor growth, invasion, migration, metastasis, chemoresistance, and *et al* in a variety of cancers via regulating the m^6^A of diverse downstream genes [15, 17, 66]. Glioblastoma (GBM, WHO grade IV glioma) is the most prevalent and malignant primary brain tumor, and characterized by rapid tumor growth, highly diffuse infiltration, chemoresistance, as well as poor prognosis, with the median survival of GBM patients less than 15 months after diagnosis [67]. Cui *et al* reported that METTL3 functions as a tumor suppressor to inhibit the growth and self-renewal of glioblastoma stem cell [47]. Consistently, Zhang *et al* reported demethylase ALKBH5 is essential for glioblastoma stem cell self-renewal and proliferation [46]. Based on different GBM cell lines used by Cui *et al* and Zhang *et al*, another two groups reported that METTL3 is highly expressed in GBM cells and plays oncogenic roles in promoting the growth, migration, invasion, and radiotherapy resistance in GBM cells [44, 45]. These diverse and somewhat conflicting roles of m^6^A in GBM are mediated by different m^6^A targets, suggesting that the roles of m^6^A in GBM depends on the contexts and specific downstream m^6^A targets. Since different m^6^A sites may direct different roles of m^6^A on GBM, targeting more specific m^6^A sites may be a promising direction in GBM therapy. It is possible that the abnormal expression of *trans* regulators of m^6^A that guide the deposition of METTL3 on highly specific downstream targets may cause dysregulation of specific m^6^A sites with more converged functions in GBM. On the other hand, the gene expression of SRSF7 and METTL3 are positively correlated in majority of cancer types of The Cancer Genome Atlas (TCGA) (Figure S9), and both *SRSF7* and *PBK* showed significantly higher gene expression in multiple cancer types (Figure S10-11), suggesting that the regulatory role of SRSF7 on m^6^A may also contribute the tumorigenicities of other cancers. Elucidating the m^6^A regulators that underlie this process may provide diverse drug targets with much fewer side effects for a variety of cancers.

## Materials and methods

### Cell culture and reagents

HEK293T cells, the Normal human astrocytes (NHA, ScienCell) and Glioma cell lines, including U87MG, LN229, A172, LN18, LN428, LN443, SNB19, T98G, U118MG, U251, and U138MG were cultured in Gibco DMEM containing 10% FBS at 37 in a humidified incubator with 5% CO_2_. All cells used in this study were confirmed mycoplasma-free.

### Tissue specimens

Both paraffin-embedded normal brain and glioma specimens were collected from glioma patients diagnosed from 2001 to 2006 at the First Affiliated Hospital of Sun Yat-sen University. Written informed consent and approval was obtained from the Institutional Research Ethics Committee of Sun Yat-sen University.

### Plasmids, siRNAs and stable cell line construction

For overexpression, the full-length coding region of *SRSF7* was subcloned into the pSin-EF2 lentiviral system. For gene silencing, short-hairpin RNA (shRNA) oligos were constructed into pLKO.1 vector. The psin-EF2-SRSF7 and pLKO.1-shSRSF7#1/2 plasmids were transfected into HEK293T cells with packing plasmids pMD2.G and psPAX2 to produce lentiviruses. Glioma cell lines were infected with these lentiviruses for 48h respectively, and later treated with puromycin for 7 days at a concentration of 0.5 μg/ml to construct stable cell lines. In addition, for the plasmids used in co-IP, the Flag-tagged full-length coding regions of *SRSF7*, *METTL3*, and *WTAP* were subcloned into pcDNA3.1 vector respectively and were transfected into U87MG cells with Lipofectamine 3000 (Invitrogen). For rescue assays, the full-length coding region of *SRSF7* with synonymous point mutations (mutate AGAACTGTATGGATTGCGAGA to AGAACCGTGTGGATCGCGCGC) was inserted into pLVX-IRES-neo plasmid to avoid being targeting by shRNAs of *SRSF7*. The PBK overexpression plasmid was constructed by inserting the full-length coding region of the major isoform of PBK (RefSeq ID: NM_018492) into pCDH-CMV-MCS-EF1-Puro vector. The two synonymous point mutations, which do not change amino acids, were introduced at m^6^A sites 1041 and 1071 by mutating A to G respectively.

Moreover, three *SRSF7* siRNAs, two *METTL3* siRNAs, two *WTAP* siRNAs, and two *IGF2BP2* siRNAs were purchased from RiboBio, China. All the sequences of siRNA oligos, PCR primers, and shRNA oligos are listed in Table S3.

### Co-immunoprecipitation (Co-IP) and Western blot

Cells were lysed with 1 × E1A lysis buffer (250 mM NaCl, 50 mM HEPES, 0.1% NP-40, 5 mM EDTA, PH adjusted at 7.5), which supplemented with 1mM PMSF and 1 × protease inhibitor cocktail (Sigma-Aldrich). The lysate was sonicated on ice and centrifuged at 4 for 15 minutes, then immunoprecipitated with Flag beads (M8823, Sigma-Aldrich) overnight.

The immunoprecipitates were washed five times with 1 × E1A lysis buffer and samples were boiled with 2 × sodium dodecyl sulphate (SDS) loading buffer at 100 for 10 minutes and ready for western blot.

Western blot was performed by using SDS-polyacrylamide gel electrophoresis, transferred onto PVDF membranes, blocked with 5% nonfat milk, and then probed with the following antibodies: anti-METTL3 (1:1000, 15073-1-AP, Proteintech), anti-METTL14 (1:1000, HPA038002, Sigma-Aldrich), anti-WTAP (1:1000, ab195380, Abcam), anti-SRSF7 (1:1000, 11044-1-AP, Proteintech), anti-PBK (1:1000, 16110-1-AP, Proteintech), anti-α-tubulin (1:1000, 66031, Proteintech), anti-Flag (1:1000, F3615, Sigma-Aldrich).

### 3D Structured Illumination Microscopy (3D-SIM)

For protein colocalization between SRSF7 and methyltransferase complex, 1.5 × 10^3^ cells of SRSF7 (Flag-tagged) overexpressed U87MG stable cell line were seeded into a chambered cover glass (Lab-Tek, Cat #155411), and the immunofluorescence staining was performed with Immunofluorescence Application Solutions Kit (CST, #12727) according to the manufacturer’s protocol. In brief, cells were fixed with 4% formaldehyde the next day, and then permeabilized with 0.2% Triton X-100 and blocked with Immunofluorescence Blocking Buffer for 1 hour, then incubated with primary antibodies (anti-METTL3: 1:1000, ab195352, Abcam; anti-METTL14: 1:200, HPA038002, Sigma-Aldrich; anti-WTAP: 1:500, ab195380, Abcam; anti-Flag: 1:200, F3165, Sigma-Aldrich) at 4 overnight. The samples were washed three times with 1× Wash Buffer the next day and probed with Alexa 488- and 647- conjugated secondary antibodies (Thermo Fisher Scientific). The images were taken by using 100× oil-immersion objective of A1R N-SIM N-STORM microscope (Nikon). All SIM images were cropped and processed with NIS Elements software.

For nuclear speckle localization of methyltransferase, the U87MG cells were transfected with *SRSF7* siRNA and negative control siRNA for 48 hours, and the immunofluorescence staining was performed as described above, and incubated with primary antibodies (anti-METTL3: 1:1000, ab195352, Abcam; anti-METTL14: 1:200, HPA038002, Sigma-Aldrich; anti-WTAP: 1:500, ab195380, Abcam; anti-SC35: 1:200, ab11826, Abcam) at 4 overnight.

### RNA isolation, quantitative reverse transcriptase PCR

Total RNA was extracted using Trizol reagent (Thermo Fisher Scientific). 1 μg RNA was reverse transcribed using GoScript Reverse Transcription Mix (A2790, Promega,) according to the manufacturer’s protocol. Quantitative real time PCR was performed using SYBR qPCR master Mix (Vazyme). Primers used in the qRT-PCR are listed in Table S3.

### m^6^A-seq

Low input m^6^A-seq was performed by using a protocol reported by Zeng *et al* [68] with some modifications. Briefly, total RNA was isolated from control U87MG cells and U87MG cells transfected with si-*SRSF7*-1 for 48 hours. A total volume of 8-10 μg total RNA was fragmented using the 10 × RNA Fragmentation Buffer (100mM Tris-HCl, 100 mM ZnCl_2_). The fragmented RNA was immunoprecipitated with 5 μg anti- m^6^A antibody (202003, Synaptic Systems), 30 μl protein-A/G magnetic beads (10002D/10004D, Thermo Fisher Scientific), 200U RNase inhibitor (N2611, Promega) in 500 μl IP Buffer (150 mM NaCl, 10 mM Tris-HCl, pH 7.5, 0.1% IGEPAL CA-630 in nuclease free H2O) at 4[for 6 hours. Then washed twice using IP buffer and eluted by competition with m^6^A sodium salt (M2780, Sigma-Aldrich). For high-throughput sequencing, both input and IP samples were used for library construction with the SMARTer Stranded Total RNA-seq Kit v2 (634413, Takara), and sequenced by Illumina HiSeq X Ten to produce 150 bp paired-end reads.

### iCLIP-seq

iCLIP was performed based on a protocol described by Yao *et al* [69] with minor modifications. Briefly, U87MG cells were UV-crosslinked with 400 mJ/cm^2^ at 254 nm and lysed with 500 μl cell lysis buffer (50 mM Tris–HCl, pH 7.4; 100 mM NaCl; 1 % NP-40; 0.1% SDS; 0.5% sodium deoxycholate), followed by immunoprecipitation with 10 μg anti-SRSF7 antibody (RN079PW, MBL), 100 μl protein A beads (10002D, Thermo Fisher Scientific) at 4 overnight and washing as described. After dephosphorylation of the 5’ ends of RNAs, linker ligation, RNA 5’ end labeling, SDS-PAGE and membrane transfer, the RNA was harvested and reverse transcribed by Superscript III (Thermo Fisher Scientific). The cDNA libraries were generated as protocol described and sequenced by Illumina NovaSeq 6000 to produce 50bp single-end reads.

### Validation of differentially methylated m^6^A sites

We used SELECT method to validate the differentially methylated m^6^A sites according to the described protocol [41]. Briefly, total RNA was mixed with 40 nM up/down primer and 5 μM dNTP in 17 μl 1 × CutSmart buffer. The mixture was annealed at a temperature gradient: 90°C, 1min; 80°C, 1min; 70°C, 1min; 60°C, 1min; 50°C, 1min, and 40°C, 6min. Then 0.5 U SplintR ligase, 0.01 U *Bst* 2.0 DNA polymerase and 10 nmol ATP was added to a final l and incubated at 40 for 20 minutes, denatured at 80 for 20 minutes, followed by qPCR. The Ct values of SELECT samples at indicated m^6^A site were normalized to the Ct values of corresponding non-modification control site. Primers used in the SELECT assay are listed in Table S3.

### Tethering assay

The full-length coding regions of *SRSF7* and *METTL3* fused with a lambda peptide sequence were cloned into pcDNA3.1, the plasmid with only a lambda peptide sequence was used as negative control. The reporter plasmid (pmirGLO-dual luciferase-5BoxB) and the effector plasmids (λ, SRSF7-λ, METTL3-λ) was transfected in U87MG cells at the ratio 1:9. The transfected cells were harvested at 24 hours after transfection and the total RNA was extracted with Trizol reagent (Thermo Fisher Scientific) and subjected to SELCET analysis [41].

Primers designed for plasmid construction and SELECT are listed in Table S3.

### RNA immunoprecipitation-qPCR analysis

Cells were harvested and lysed in NP-40 lysis buffer (20 mM Tris–HCl at pH 7.5, 100 mM KCl, 5 mM MgCl_2_, and 0.5% NP-40), then cell lysates were immunoprecipitated with 10 μg anti-METTL3 (15073-1-AP, Proteintech), or anti-METTL14 (26158-1-AP, Proteintech), or anti-WTAP (ab195380, Abcam) respectively, and 100 μl protein G beads (10004D, Thermo Fisher Scientific) at 4[overnight, followed by DNase I treatment, proteinase K treatment.

The bound RNAs were extracted by Trizol reagent, reverse transcribed into cDNAs, and subjected to qPCR analysis.

### Cell proliferation, colony formation assay, migration assay, and sphere formation assay

For cell growth curve, 1 × 10^3^ cells were seeded into 96-well plates and stained with MTT (Sigma-Aldrich) dye, and measured the absorbance at 570 nm. Colony formation was performed by seeding cells (1 × 10^3^) into 12-well plates, cultured for 7 days, then fixed with methanol and stained with Crystal violet.

For EdU assays, 2 × 10^4^ cells were seeded into 48-well plates and EdU assays were performed using the EdU Cell Proliferation Assay Kit (Cat.C10310-1, RiboBio, China). Cell migration assays were performed by seeding 2 × 10^4^ cells into 24-well transwell polycarbonate membrane cell culture inserts and stained with Crystal violet.

For sphere formation assay, 3 × 10^3^ cells were seeded into Ultra-Low Attachment Multiple Well Plate, and cultured in the stem cell culture condition for 7 days.

### Intracranial Xenograft

Five-week-old female BALB/c nude mice was obtained from Beijing Vital River (Beijing, China) and divided into two groups (SRSF7-KD and control, n=6 per group). Each mouse was injected with 5 × 10^5^ U87MG cells which expressing luciferase in the right cerebrum. Tumor growth was monitored by Bioluminescent imaging every week. The study protocol was approved by the Institutional Animal Care and Use Committee of Sun Yat-sen University Cancer Center.

### RNA stability assays

Cells were treated with 5 μg/ml actinomycin D (A9415, Sigma-Aldrich) and collected at 0 hour, 3 hours, 6 hours, 9 hours after treatment. The total RNA was isolated, reverse transcribed into cDNA, and subjected to qPCR analysis.

### m^6^A-seq data analyses

The first end of the raw paired-end reads of the m^6^A-seq were trimmed to 50 bp from the 3’ end for m^6^A peak calling and downstream analyses. We mapped the reads to hg19 human genome using HISTA2 (v2.1.0) [70]. The m^6^A peaks were identified according to the methods as described in our previous paper [14, 35], which was modified from the method published by Dominissini et al [12]. We created 100 bp sliding windows with 50 bp overlapped along the longest isoforms of each Ensembl annotated gene and calculated the RPKM (Reads Per Kilobase of transcript, per Million mapped reads) for each window for IP and input respectively. For each window, the ratio of RPKM+1 between IP and input were calculated as the intensity. The winscore of each window was then calculated as the ratio of intensities between this window and the median of all windows in the same gene. Windows with RPKM > 10 in the IP and winscore (enrichment score) > 2 were defined as the enriched windows in each sample. The m^6^A peaks were defined as the enriched windows with winscores greater than neighboring windows. The overlapped or just neighboring peaks of the two biological replicates were merged into larger windows and the 100 bp region in the middle of the merged peak were considered as the common peaks, which were further filtered by requiring the winscores > 2 in both replicates. The distributions of m^6^A peaks along 30 bins of mRNA were calculated as we have previously described [14].

The m^6^A ratio, which quantifies m^6^A peaks, of each m^6^A peak were calculated as the ratio of peak RPKM between IP and input. To calculate the fold change of m^6^A ratios upon *SRSF7* knockdown, we first took the union of the m^6^A peaks of all samples. The union peaks of two replicates were merged, centralized, and filtered to obtain a set of 100 bp peak regions in the same way as above described for obtaining common peaks. To avoid using the unreliable m^6^A ratios due to tiny denominators, we filtered out the peaks with input window RPKM < 5 at least one sample or m^6^A ratio < 0.1 in any control samples. Then the m^6^A peaks with fold change of m^6^A ratios upon *SRSF7* knockdown > 1.5 or < 2/3 were determined as the up-regulated or down-regulated m^6^A peaks respectively. The data were visualized using the Integrative Genomics Viewer (IGV) tool [71], the biological replicates were merged and the average read coverages were used for visualization. StringTie (v1.3.4d) [72] was used to calculate the TPMs (Transcripts Per Million) of Ensembl annotated genes using the input libraries. We filtered out the genes with mean TPMs < 1 in control samples to avoid using unreliable fold change of TPMs due to tiny denominators. Differentially expressed genes were determined using DESeq2 [73] according to the read counts of genes calculated by HTSeq [74]. The genes with FDR < 0.05 and mean CPM (Counts per Million) > 100 were determined as the differentially expressed genes. Gene Ontology analysis was performed using DAVID [75] with all expressed genes (TPM >1) as background. The GSEA analysis was performed using GSEA (version 2.2.2.0) [76] based on the predefined gene sets from the Molecular Signatures Database (MSigDB v5.0) [76].

### Analyses of the clinical data of glioma patients

The gene expression, mutation, and clinical data of glioma patients were downloaded from CGGA database (http://www.cgga.org.cn/) [36] We used the Cox regression to examine the correlations between gene expression indexes of the cancer module and patient survival in each cancer type. The gene expression data of all cancer types were downloaded from TCGA (https://tcga-data.nci.nih.gov/).

### iCLIP-seq data analyses

We used the CLIP Tool Kit (CTK) to call the peaks from the iCLIP-seq data according to the described data processing procedure of iCLIP-seq [38]. HOMER software [77] was used for motif enrichment analysis with randomly permutated sequences as the background. The overlapped peaks between the peaks of m^6^A-seq and iCLIP-seq were determined as the peaks with distances < 100 bp using BEDTools [78].

### Alternative splicing and alternative polyadenylation (APA) analyses

We used rMATS [57] to perform the differential alternative splicing analysis using the input RNAs of m^6^A-seq with FDR < 0.05 as the threshold of significance. The binding enriched of SRSF7 around splicing events were analyzed using rMAPS2. To test whether the genes with alternative splicing and the genes with SRSF7 regulated m^6^A are significantly overlapped, we only considered all m^6^A modified genes with rMATS detected alternative splicing in the Chi-square test. Differential alternative polyadenylation (APA) analysis was performed using DaPars [59] with FDR < 0.1 as the threshold of significance.

### Statistics

Comparisons between two groups were performed using Student’s two-tailed *t* test. Comparisons during more than two groups are performed using ANOVA. Data represent mean ± SEM, *P* value or adjusted *P* value for ANOVA less than 0.05 were considered statistically significant. Survival curves were plotted by the Kaplan–Meier method and compared by the log-rank test. The statistics of bioinformatic analyses were all described along with the results or figures.

### Data Availability Statement

The raw sequencing reads of m^6^A-seq and iCLIP-seq have been deposited in Genome Sequence Archive (GSA) for Human under the accession code HRA001166 (reviewer access link: https://ngdc.cncb.ac.cn/gsa-human/s/vrR56Et5).

## Authors’ contributions

JW and JLi conceived and supervised the project; YC, CX, JLan, WL performed experiments with the assistances from CY, XL, XH, XS, YH, ZL, SZ, GW, MY, MT, RY, XL, GG, WZ; SA and WC performed bioinformatics analyses with the help of ZW; YC draft the manuscript; JW and JLi revised the manuscript. All authors read and approved the final manuscript.

## Competing interests

The authors declare no competing interests.

## Acknowledgments

We thank Jianzhao Liu for providing the vectors of tethering assay. This work was supported by the National Key R&D Program of China [2018YFA0107200 to JW], the National Natural Science Foundation of China [81830082, 82030078, and 81621004 to JL; 31771446 and 31970594 to JW], Guangzhou Science and Technology Program [201904010181 to JW], Guangzhou Science and Technology Plan Projects [201803010098 to JL]; Natural Science Foundation of Guangdong Province [2018B030311009 to JL].

## Supplementary figure legends

**Figure S1 Interaction between SRSF7 and methyltransferase complex**

**A.** Western blots showing Flag-tagged SRSF7 interacts with endogenous METTL3, METTL14 and WTAP with RNase treatment in 293T cells. **B.** Schematic diagram of the truncated regions of *SRSF7.* FL: Full length, RRM: RNA recognition motif, Zn: Zinc knuckle, RS: arginine/serine.

**Figure S2 SRSF7 specifically facilitates m^6^A methylation near its binding sites**

**A.** Enriched motifs in m^6^A peaks of control and *SRSF7*-KD U87MG cells. **B.** Normalized distributions of m^6^A peaks across 5’UTR, CDS, and 3’UTR of mRNA in U87MG cells transfected with scramble (si-NC) and siRNAs of *SRSF7* (si-*SRSF7*) respectively. **C.** Box plot comparing the m^6^A ratios of the m^6^A peaks in control and *SRSF7*-KD U87MG cells. **D.** Heatmap representing the Z-score transformed m^6^A ratios in si-NC and si-*SRSF7* in U87MG cells respectively. **E, F.** GO (E) and KEGG (F) enrichment analyses of genes with down-regulated m^6^A peaks upon *SRSF7* knockdown. **G.** Pie chart showing the fractions of SRSF7 iCLIP-seq peaks located in different regions of genes. **H.** Normalized distributions of SRSF7 iCLIP-seq peaks colocalized with m^6^A peaks across 5’UTR, CDS, and 3’UTR of mRNA in U87MG cells. **I.** Plot of cumulative fraction of log_2_ fold change of m^6^A ratios upon *SRSF7* knockdown using si-*SRSF7* for the m^6^A peaks within the orange module and all other modules respectively. *P* value of two-tailed Wilcoxon test is indicated.

**Figure S3 SRSF7 regulates gene expression**

**A.** Heatmap representing the Z-score transformed gene expression of differentially expressed genes between control and SRSF7-KD U87MG cells. **B, C.** GO enrichment analyses of genes with down-regulated (B) and up-regulated (C) gene expression upon *SRSF7* knockdown. **D-F.** GSEA plot for the gene expression changes due to *SRSF7* knockdown in U87MG cells.

**Figure S4 SRSF7 directly targets and facilitates the methylation of m^6^A on genes involved in cell proliferation and migration**

**A.** KEGG enrichment of the corresponding genes with the overlapped m^6^A peaks between down-regulated m^6^A peaks upon *SRSF7* knockdown and SRSF7 iCLIP-seq peaks. **B.** GO enrichment analysis of genes with SRSF7 iCLIP-seq peaks not colocalize with m^6^A peaks. **C.** Tracks displaying the read coverage of IPs and inputs of m^6^A-seq as well as the SRSF7 iCLIP-seq on *ROBO1*. The SRSF7 directly regulated m^6^A peak is highlighted.

**Figure S5 SRSF7 promotes the migration and proliferation of GBM cells**

**A.** Representative images of colony formation assay in U87MG and LN229 cells overexpressed SRSF7. **B.** Western blot showing efficiently knockdown of SRSF7 in U87MG and LN229 cells transduced with control shRNA or *SRSF7* shRNA respectively. **C.** Representative images of transwell migration assay in U87MG and LN229 cells transduced with control shRNA or *SRSF7* shRNA respectively, Scar bars: 50 μm. **D.** Representative images of EdU staining assays and bar plot comparing the EdU positive rates in U87MG and LN229 cells transduced with control shRNA or *SRSF7* shRNA respectively. Data are presented as mean ± SEM, n = 5. *** *P* < 0.001. One-way ANOVA with Dunnett’s post hoc μm. **E.** Western blot showing the protein level of SRSF7 in U87MG and LN229 cells transduced with sh*SRSF7* together with empty vector and *SRSF7* with synonymous mutations. **F.** Representative images of colony formation assay in U87MG and LN229 cells with control, SRSF7 knockdown, and SRSF7 knockdown rescued by SRSF7 overexpression. **G.** Representative images of EdU staining assays and bar plot comparing the EdU positive rates in U87MG and LN229 cells with control, SRSF7 knockdown, and SRSF7 knockdown rescued by SRSF7 overexpressed. Data are presented as mean ± SEM, n = 5. * *P* < 0.05, ** *P* < 0.01, *** *P* < 0.001. One-way ANOVA with Tukey’s post hoc test. Scar bars: 50 μm. **H.** Sphere formation results in U87MG cells upon SRSF7 depletion and overexpressed. Scar bars: 100 μm.

**Figure S6 SRSF7 promotes the proliferation and migration of glioblastoma cells partially dependent on METTL3**

**A, B.** Gene expression change of *METTL3*, *METTL14*, and *WTAP* in U87MG cells (A) and (B) transfected with scramble (si-NC) and siRNA of *SRSF7* (si-*SRSF7-*1, si-*SRSF7*-2 and si-*SRSF7*-3) respectively. Data are presented as mean ± SEM, n = 3. *** *P* < 0.001. ns: no significant difference. One-way ANOVA with Dunnett’s post hoc test. **C-E.** Western blot showing the protein level of METTL3, METTL14, WTAP, and SRSF7 upon *SRSF7* knockdown (C-D) or *SRSF7* overexpression (E) in U87MG and LN229 cells. **F.** Western blot showing the protein level of METTL3 and SRSF7 in U87MG cells transfected with si-NC and si-METTL3. **G.** Western blot showing the protein level of WTAP and SRSF7 in U87MG cells transfected with si-NC and si-WTAP. **H-J.** 3D-SIM imaging of colocalization of METTL3 (H), METTL14 (I), and WTAP (J) with the nuclear speckle marker SC35. Scale Bar: 2 μm.

**Figure S7 SRSF7 promotes the proliferation and migration of GBM cells partially through increasing the stability of *PBK* mRNA**

**A.** Kaplan-Meier survival analysis based on *PBK* expression in GBM patients from CGGA dataset. **B.** Relative mRNA expression level of *PBK* in GBM patients from CGGA dataset. **C.** Scatter plot showing the correlation between *METTL3* and *PBK* gene expression across GBM patients from CGGA dataset, the *P* value and correlation coefficient are indicated. **D.** Western blot showing the protein level of PBK and SRSF7 in U87MG and LN229 cells upon *SRSF7* knockdown and rescued by co-transducing full-length WT *PBK* CDS regions. **E.** Colony formation results in U87MG and LN229 cells upon *SRSF7* knockdown and rescued by co-transducing full-length WT *PBK* CDS regions. **F, G.** Bar plot showing the relative mRNA level of *SRSF7* (F) and *METTL3* (G) in SRSF7 overexpressed U87MG cells transfected without or with si-*METTL3*-1 as indicated. Data are presented as mean ± SEM, n = 3. *** *P* < 0.001. One-way ANOVA with Tukey’s post hoc test. **H.** Relative mRNA expression level of *PBK* in U87MG cells transfected with scramble (si-NC) and siRNA of *IGF2BP2* (si-*IGF2BP2*-1, si-*IGF2BP2*-2) respectively. Data are presented as mean ± SEM, n = 3. ** *P* < 0.01, *** *P* < 0.001. One-way ANOVA with Dunnett’s post hoc test. **I.** Scatter plot showing the correlation between *IGF2BP2* and *PBK* gene expression across GBM patients from CGGA dataset, the *P* value and correlation coefficient are indicated.

**Figure S8: Motif analyses of up-regulated m^6^A peaks and SRSF7 iCLIP-seq peaks that affect m^6^A**

**A.** Motifs enriched in the up-regulated m^6^A peaks using all m^6^A peaks as background. **B.** Motifs enriched in the SRSF7 iCLIP-seq peaks that affect m^6^A using all iCLIP-seq peaks as background.

**Figure S9: Scatter plot showing the correlation coefficiences (r) and -log_10_P value of the gene expression between *SRSF7* and *METTL3* in multiple cancers of TCGA dataset**

**Figure S10: Boxplots comparing the gene expression of *SRSF7* in cancer (red) and normal (grey) for multiple cancers of TCGA dataset. *: P < 0.01.**

**Figure S11: Boxplots comparing the gene expression of *PBK* in cancer (red) and normal (grey) for multiple cancers of TCGA dataset. *: P < 0.01**

**Table S1: Information of SRSF7 iCLIP-seq peaks in U87MG cells**

**Table S2: Information of SRSF7 directly regulated m6A peaks**

**Table S3: Sequences of primers, siRNAs and shRNA**

## Notes

http://www.cgga.org.cn/

https://tcga-data.nci.nih.gov/

### Competing Interest Statement

The authors have declared no competing interest.

