## Supplementary Figures for "Serine/arginine-rich splicing factor 7 plays oncogenic roles through specific regulation of m^6^A RNA modification"

**Figure S1 Interaction between SRSF7 and methyltransferase complex**
**A.** Western blots showing Flag-tagged SRSF7 interacts with endogenous METTL3, METTL14 and WTAP with RNase treatment in 293T cells. **B.** Schematic diagram of the truncated regions of *SRSF7.* FL: Full length, RRM: RNA recognition motif, Zn: Zinc knuckle, RS: arginine/serine.

A

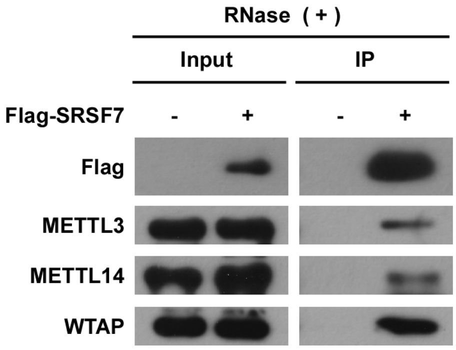


B

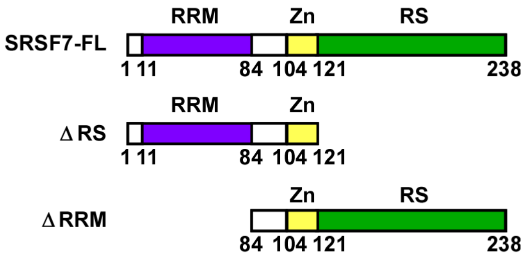

**Figure S2 SRSF7 specifically facilitates m$^6$A methylation near its binding sites**
**A.** Enriched motifs in m$^6$A peaks of control and *SRSF7*-KD U87MG cells. **B.** Normalized distributions of m$^6$A peaks across 5'UTR, CDS, and 3'UTR of mRNA in U87MG cells transfected with scramble (si-NC) and siRNAs of *SRSF7* (si-*SRSF7*) respectively. **C.** Box plot comparing the m$^6$A ratios of the m$^6$A peaks in control and *SRSF7*-KD U87MG cells. **D.** Heatmap representing the Z-score transformed m$^6$A ratios in si-NC and si-*SRSF7* in U87MG cells respectively. **E, F.** GO (E) and KEGG (F) enrichment analyses of genes with down-regulated m$^6$A peaks upon *SRSF7* knockdown. **G.** Pie chart showing the fractions of SRSF7 iCLIP-seq peaks located in different regions of genes. **H.** Normalized distributions of SRSF7 iCLIP-seq peaks colocalized with m$^6$A peaks across 5'UTR, CDS, and 3'UTR of mRNA in U87MG cells. **I.** Plot of cumulative fraction of $\log_2$ fold change of m$^6$A ratios upon *SRSF7* knockdown using si-*SRSF7* for the m$^6$A peaks within the orange module and all other modules respectively. *P* value of two-tailed Wilcoxon test is indicated.

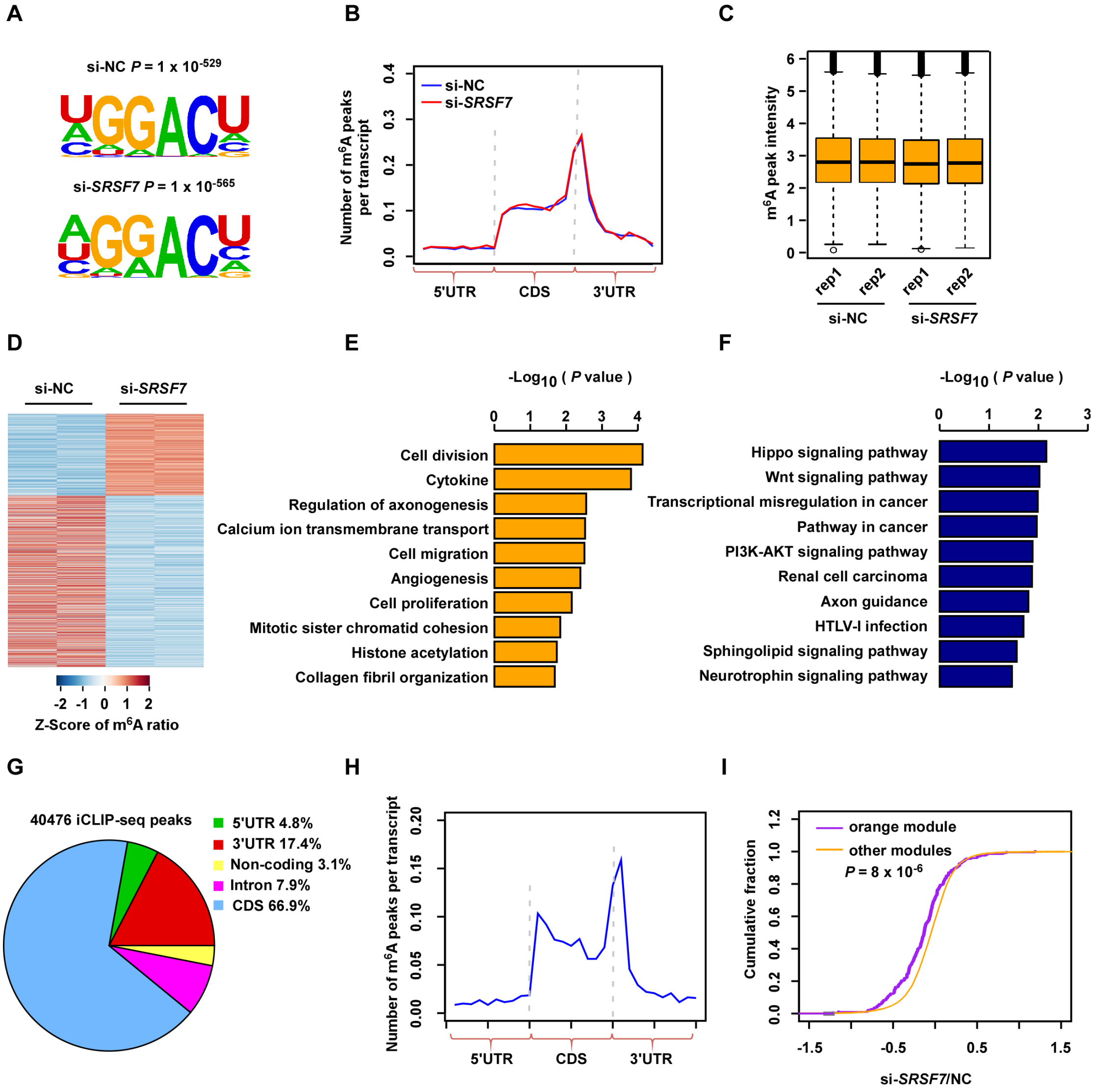

**Figure S3 SRSF7 regulates gene expression**

**A.** Heatmap representing the Z-score transformed gene expression of differentially expressed genes between control and SRSF7-KD U87MG cells. **B, C.** GO enrichment analyses of genes with down-regulated (B) and up-regulated (C) gene expression upon *SRSF7* knockdown. **D-F.** GSEA plot for the gene expression changes due to *SRSF7* knockdown in U87MG cells.

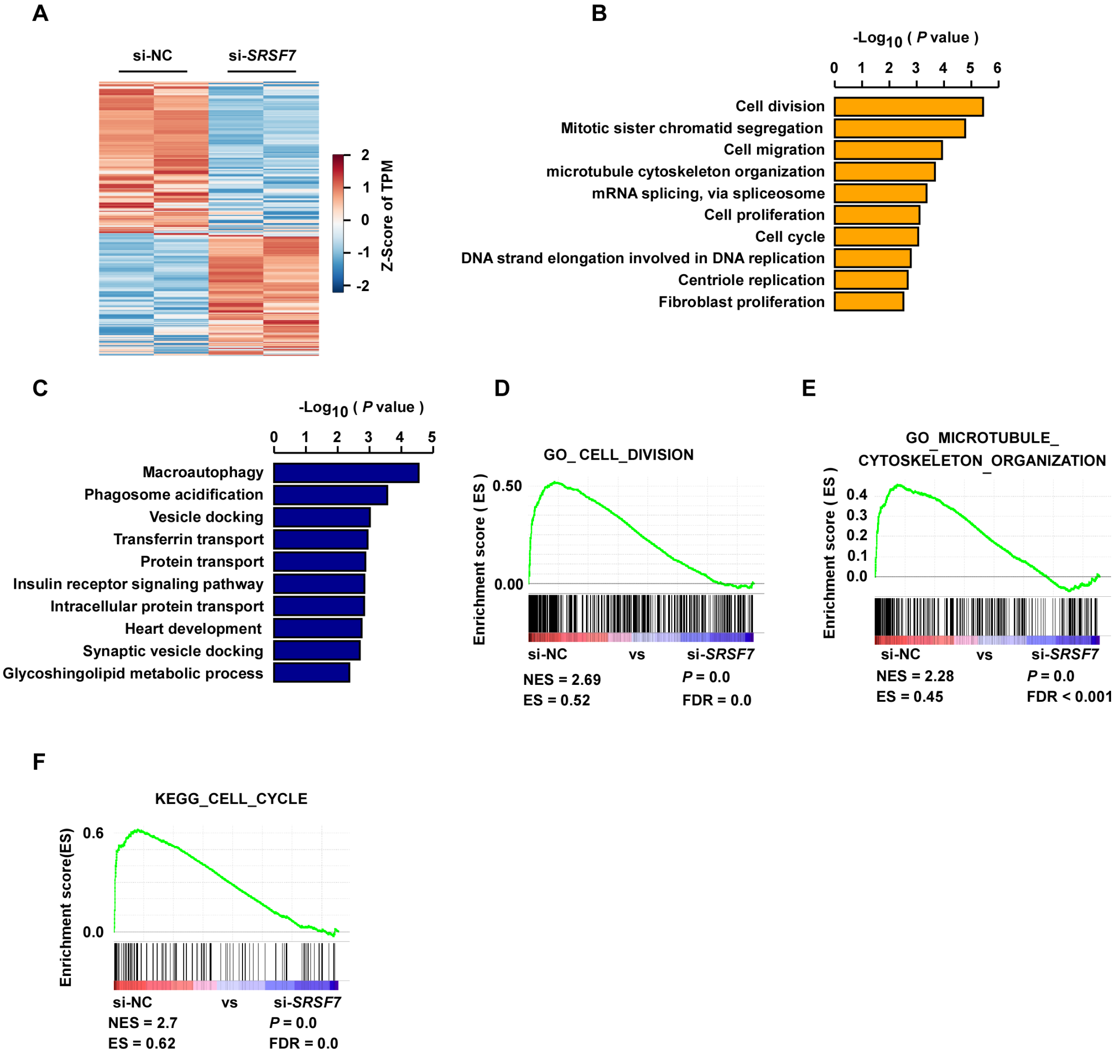

**Figure S4 SRSF7 directly targets and facilitates the methylation of m$^6$A on genes involved in cell proliferation and migration**

**A.** KEGG enrichment of the corresponding genes with the overlapped m$^6$A peaks between down-regulated m$^6$A peaks upon *SRSF7* knockdown and SRSF7 iCLIP-seq peaks. **B.** GO enrichment analysis of genes with SRSF7 iCLIP-seq peaks not colocalize with m$^6$A peaks. **C.** Tracks displaying the read coverage of IPs and inputs of m$^6$A-seq as well as the SRSF7 iCLIP-seq on *ROBO1*. The SRSF7 directly regulated m$^6$A peak is highlighted.

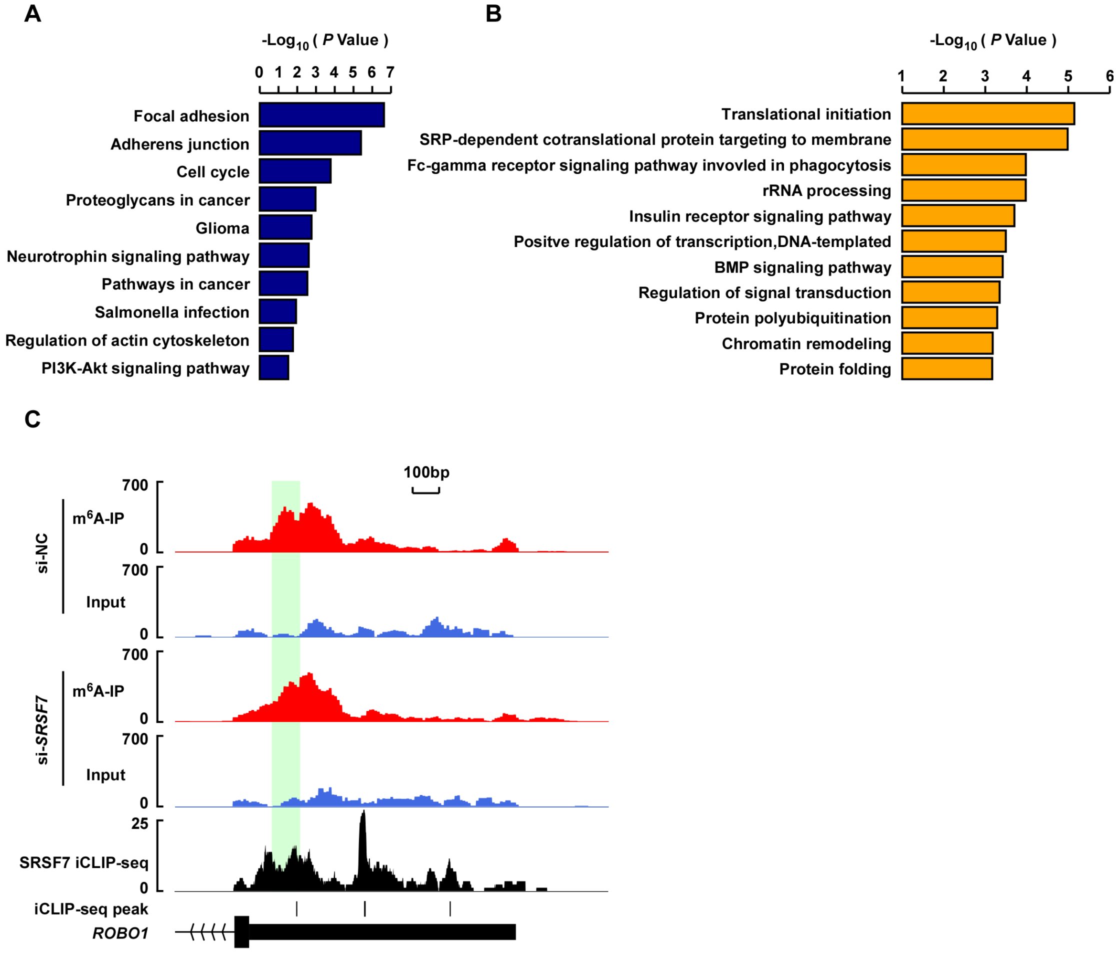

**Figure S5 SRSF7 promotes the migration and proliferation of GBM cells**

**A.** Representative images of colony formation assay in U87MG and LN229 cells overexpressed SRSF7. **B.** Western blot showing efficiently knockdown of SRSF7 in U87MG and LN229 cells transduced with control shRNA or *SRSF7* shRNA respectively. **C.** Representative images of transwell migration assay in U87MG and LN229 cells transduced with control shRNA or *SRSF7* shRNA respectively, Scar bars: 50 μm. **D.** Representative images of EdU staining assays and bar plot comparing the EdU positive rates in U87MG and LN229 cells transduced with control shRNA or *SRSF7* shRNA respectively. Data are presented as mean ± SEM, n = 5. *** $P < 0.001$. One-way ANOVA with Dunnett's post hoc test. Scar bars: 50 μm. **E.** Western blot showing the protein level of SRSF7 in U87MG and LN229 cells transduced with sh*SRSF7* together with empty vector and *SRSF7* with synonymous mutations. **F.** Representative images of colony formation assay in U87MG and LN229 cells with control, SRSF7 knockdown, and SRSF7 knockdown rescued by SRSF7 overexpression. **G.** Representative images of EdU staining assays and bar plot comparing the EdU positive rates in U87MG and LN229 cells with control, SRSF7 knockdown, and SRSF7 knockdown rescued by SRSF7 overexpressed. Data are presented as mean ± SEM, n = 5. * $P < 0.05$, ** $P < 0.01$, *** $P < 0.001$. One-way ANOVA with Tukey's post hoc test. Scar bars: 50 μm. **H.** Sphere formation results in U87MG cells upon SRSF7 depletion and overexpressed. Scar bars: 100 μm.

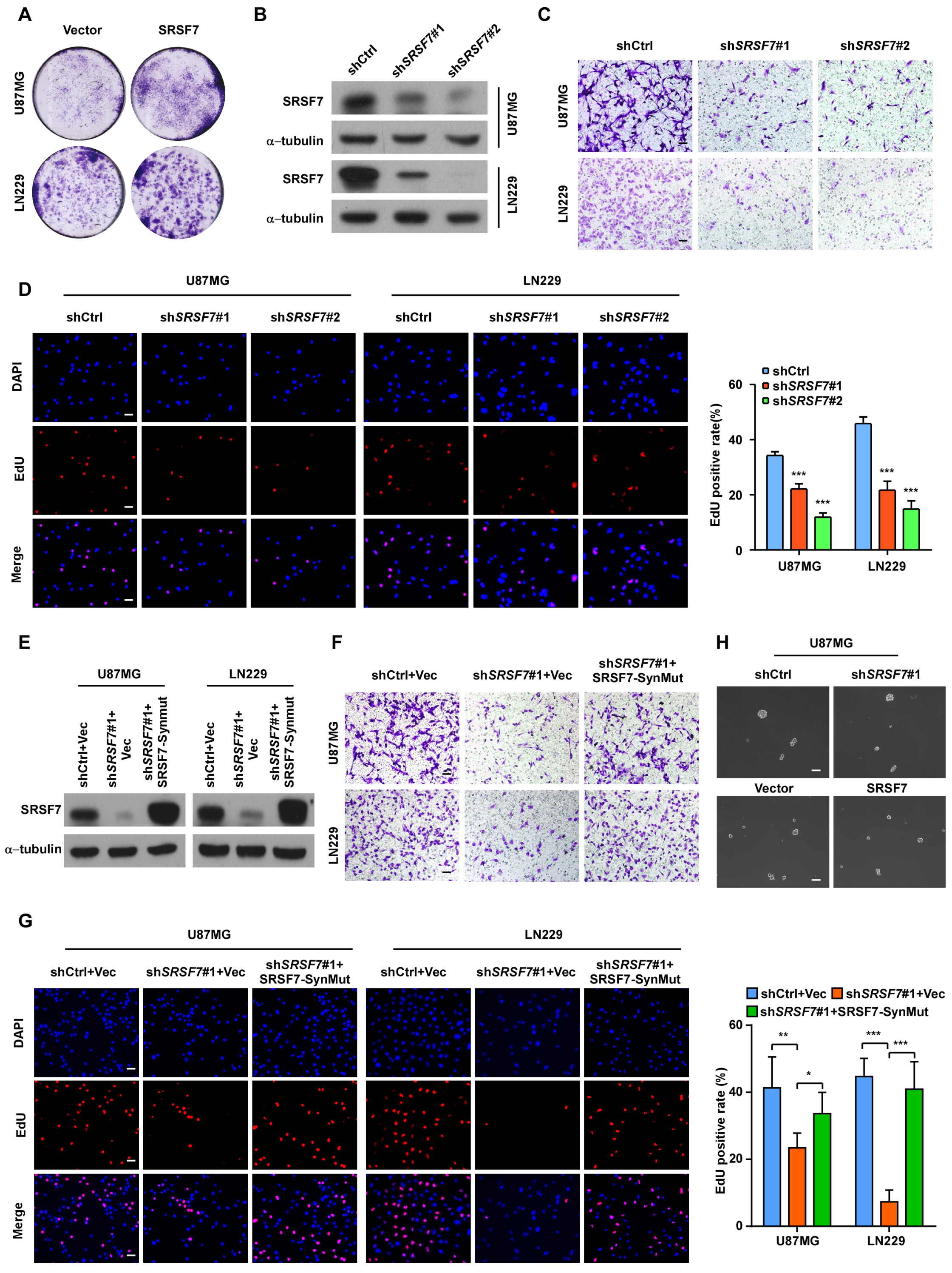
A
Vector
SRSF7
U87MG
LN229
B
shCtrl
shSRSF7#1
shSRSF7#2
SRSF7
α–tubulin
U87MG
LN229
C
shCtrl
shSRSF7#1
shSRSF7#2
D
U87MG
LN229
DAPI
EdU
Merge
shCtrl
shSRSF7#1
shSRSF7#2
EdU positive rate(%)
60
40
20
0
***
E
shCtrl+Vec
shSRSF7#1+Vec
shSRSF7#1+SRSF7-Synmut
F
shCtrl+Vec
shSRSF7#1+Vec
shSRSF7#1+SRSF7-SynMut
H
Vector
SRSF7
G
shCtrl+Vec
shSRSF7#1+Vec
shSRSF7#1+SRSF7-SynMut
EdU positive rate (%)
**
*
***

**Figure S6 SRSF7 promotes the proliferation and migration of glioblastoma cells partially dependent on METTL3**

**A, B.** Gene expression change of *METTL3*, *METTL14*, and *WTAP* in U87MG cells (A) and (B) transfected with scramble (si-NC) and siRNA of *SRSF7* (si-*SRSF7*-1, si-*SRSF7*-2 and si-*SRSF7*-3) respectively. Data are presented as mean ± SEM, n = 3. *** $P < 0.001$. ns: no significant difference. One-way ANOVA with Dunnett's post hoc test. **C-E.** Western blot showing the protein level of METTL3, METTL14, WTAP, and SRSF7 upon *SRSF7* knockdown (C-D) or *SRSF7* overexpression (E) in U87MG and LN229 cells. **F.** Western blot showing the protein level of METTL3 and SRSF7 in U87MG cells transfected with si-NC and si-METTL3. **G.** Western blot showing the protein level of WTAP and SRSF7 in U87MG cells transfected with si-NC and si-WTAP. **H-J.** 3D-SIM imaging of colocalization of METTL3 (H), METTL14 (I), and WTAP (J) with the nuclear speckle marker SC35. Scale Bar: 2 µm.

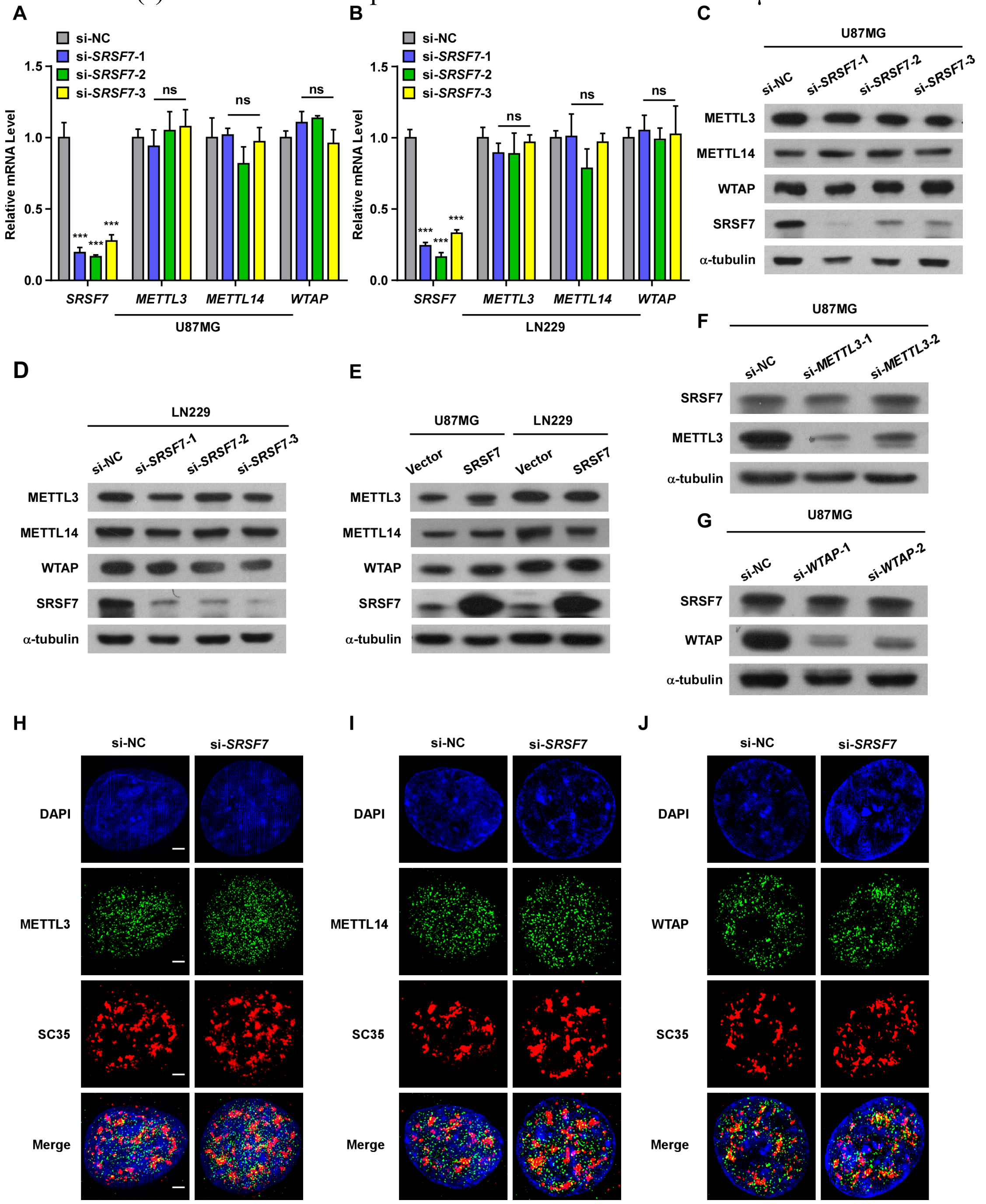

**Figure S7 SRSF7 promotes the proliferation and migration of GBM cells partially through increasing the stability of *PBK* mRNA**

**A.** Kaplan-Meier survival analysis based on *PBK* expression in GBM patients from CGGA dataset. **B.** Relative mRNA expression level of *PBK* in GBM patients from CGGA dataset. **C.** Scatter plot showing the correlation between *METTL3* and *PBK* gene expression across GBM patients from CGGA dataset, the *P* value and correlation coefficient are indicated. **D.** Western blot showing the protein level of PBK and SRSF7 in U87MG and LN229 cells upon *SRSF7* knockdown and rescued by co-transducing full-length WT *PBK* CDS regions. **E.** Colony formation results in U87MG and LN229 cells upon *SRSF7* knockdown and rescued by co-transducing full-length WT *PBK* CDS regions. **F, G.** Bar plot showing the relative mRNA level of *SRSF7* (F) and *METTL3* (G) in SRSF7 overexpressed U87MG cells transfected without or with si-*METTL3*-1 as indicated. Data are presented as mean ± SEM, n = 3. *** $P < 0.001$. One-way ANOVA with Tukey's post hoc test. **H.** Relative mRNA expression level of *PBK* in U87MG cells transfected with scramble (si-NC) and siRNA of *IGF2BP2* (si-*IGF2BP2*-1, si-*IGF2BP2*-2) respectively. Data are presented as mean ± SEM, n = 3. ** $P < 0.01$, *** $P < 0.001$. One-way ANOVA with Dunnett's post hoc test. **I.** Scatter plot showing the correlation between *IGF2BP2* and *PBK* gene expression across GBM patients from CGGA dataset, the *P* value and correlation coefficient are indicated.

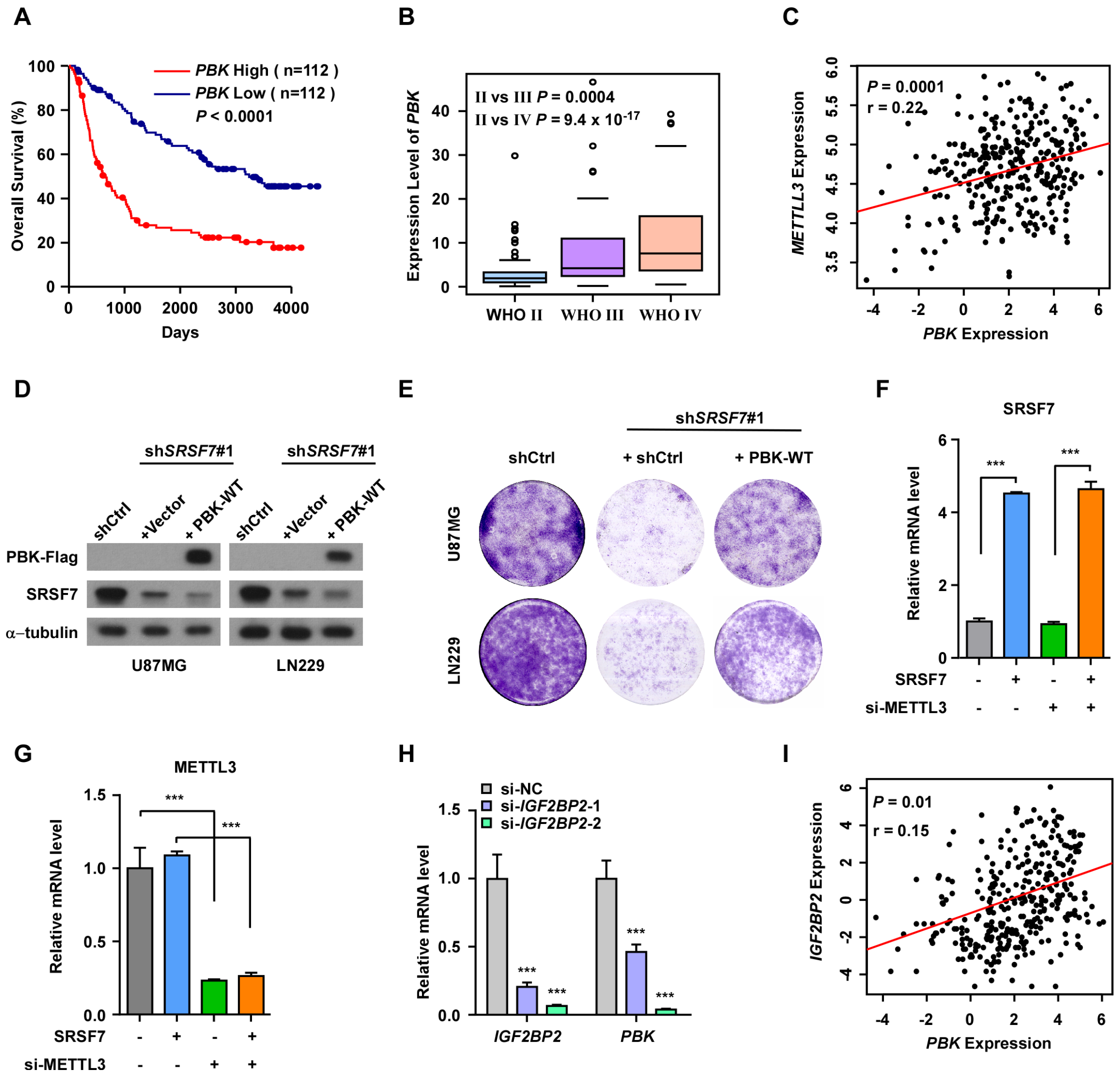

**Figure S8: Motif analyses of up-regulated m$^6$A peaks and SRSF7 iCLIP-seq peaks that affect m$^6$A.**

**A.** Motifs enriched in the up-regulated m$^6$A peaks using all m$^6$A peaks as background.

**B.** Motifs enriched in the SRSF7 iCLIP-seq peaks that affect m$^6$A using all iCLIP-seq peaks as background.

**A**

| Rank | motif | *P* value |
|---|---|---|
| 1 | 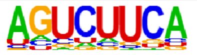 | 1 x $10^{-6}$ |
| 2 | 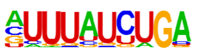 | 1 x $10^{-6}$ |
| 3 | 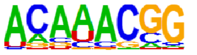 | 1 x $10^{-6}$ |
| 4 | 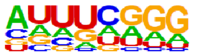 | 1 x $10^{-5}$ |
| 5 | 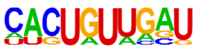 | 1 x $10^{-5}$ |
| 6 | 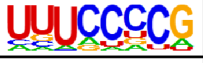 | 1 x $10^{-5}$ |
| 7 | 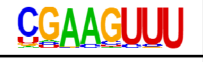 | 1 x $10^{-4}$ |
| 8 | 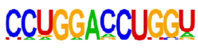 | 1 x $10^{-4}$ |

**B**

| Rank | motif | *P* value |
|---|---|---|
| 1 | 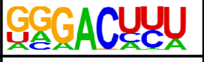 | 1 x $10^{-9}$ |
| 2 | 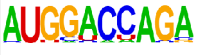 | 1 x $10^{-9}$ |
| 3 | 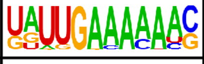 | 1 x $10^{-9}$ |
| 4 | 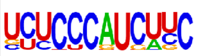 | 1 x $10^{-8}$ |
| 5 | 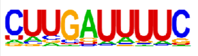 | 1 x $10^{-8}$ |
| 6 | 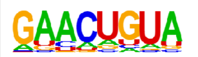 | 1 x $10^{-8}$ |
| 7 | 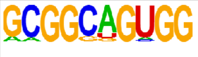 | 1 x $10^{-8}$ |
| 8 | 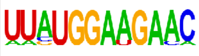 | 1 x $10^{-6}$ |
| 9 | 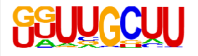 | 1 x $10^{-6}$ |
| 10 | 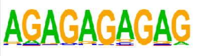 | 1 x $10^{-4}$ |

**Figure S9: Scatter plot showing the correlation coefficiences (r) and -log$_{10}$P value of the gene expression between *SRSF7* and *METTL3* in multiple cancers of TCGA dataset**

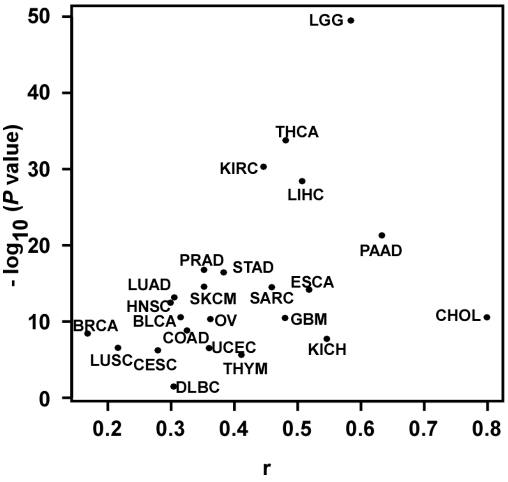

**Figure S10: Boxplots comparing the gene expression of *SRSF7* in cancer (red) and normal (grey) for multiple cancers of TCGA dataset. *: P < 0.01**

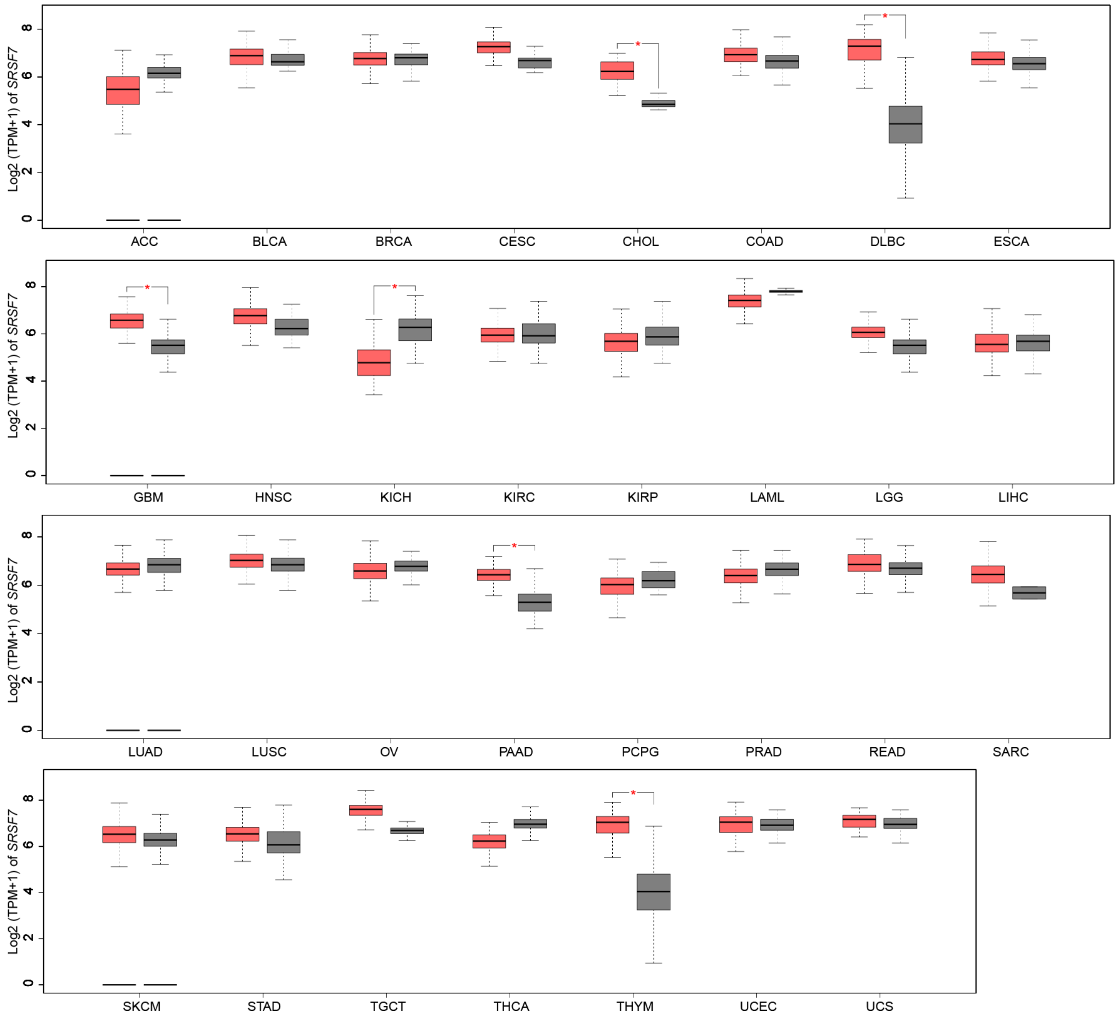

**Figure S11: Boxplots comparing the gene expression of *PBK* in cancer (red) and normal (grey) for multiple cancers of TCGA dataset. *: P < 0.01**

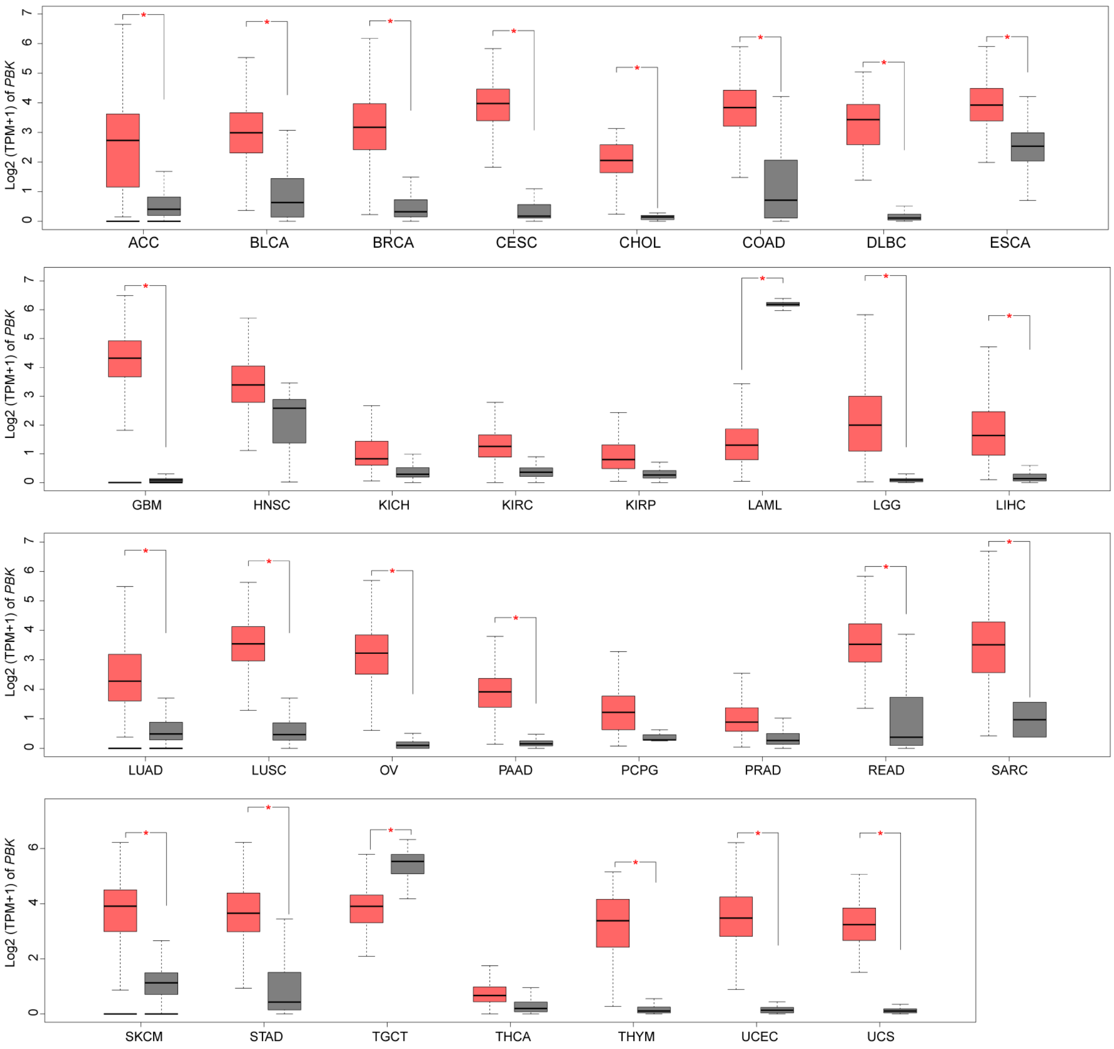
